# Scrutinizing science to save lives: uncovering flaws in the data linking L-type calcium channels blockers to CRAC channels and heart failure

**DOI:** 10.1101/2024.02.06.579229

**Authors:** Gary S. Bird, Yu-Ping Lin, Charles J. Tucker, Geoffrey Mueller, Min Shi, Sandosh Padmanabhan, Anant B. Parekh

## Abstract

Hypertension is estimated to affect almost 1 billion people globally and significantly increases risk of myocardial infarction, heart failure, stroke, retinopathy and kidney disease. One major front line therapy that has been used for over 50 years involves L-type Ca^2+^ channel blockers (LCCBs). One class of LCCBs is the dihydropyridine family, with amlodipine being widely prescribed regardless of gender, race, ethnicity or age. In 2020, Johnson et al.^7^ reported that all LCCBs significantly increased the risk of heart failure, and attributed this effect to non-canonical activation of store-operated Ca^2+^ entry. A major approach on which they based many of their arguments was to measure cytosolic Ca^2+^ using the fluorescent Ca^2+^ indicator dye fura-2. We recently demonstrated that amlodipine is highly fluorescent within cells and overwhelms the fura-2 signal, precluding the use of the indicator dye with amlodipine^24^. Our meta-analyses and prospective real world study showed that dihydropyridines were not associated with an increase in heart failure, likely explained by the lack of consideration by Johnson et al.^7^ of well-known confounding factors such as age, race, obesity, prior anti-hypertensive treatment or diabetes^24^. Trebak and colleagues have responded to our paper with a forthright and unwavering defence of their work^27^. In this paper, we carry out a forensic dissection of Johnson et al.,^7^ and conduct new experiments that address directly points raised by Trebak et al. ^27^. We show that there are major flaws in the design and interpretation of their key experiments, that fura-2 cannot be used with amlodipine, that there are fundamental mathematical misunderstandings and mistakes throughout their study leading to critical calculations on heart failure that are demonstrably wrong, and several of their own results are inconsistent with their interpretation. We therefore believe the study by Johnson et al. ^7^ is flawed at many levels and we stand by our conclusions.

## Introduction

Globally, around one in five adults is diagnosed with the ‘silent killer’ hypertension, which significantly increases risk of myocardial infarction, heart failure, stroke, retinopathy and kidney disease^1^. 116 million Americans are thought to have hypertension or are taking medication for its treatment^2^. The European Commission reports 1 in 5 Europeans are hypertensive^3^, the WHO estimates 46% of adults over 25 years of age in Africa have high blood pressure and between 15–35% of urban adult populations of Asia are hypertensive^4^. Several different classes of drug are used to treat hypertension, and L-type calcium channel blockers (LCCBs) are a universal first line treatment^5^. The most effective and widely prescribed LCCB is the dihydropyridine amlodipine. In India, where amlodipine is the most widely administered anti-hypertensive agent^6^, the drug costs less than $3 for an entire month’s supply, making it affordable to low-income families. Challenging the safety of amlodipine and other LCCBs therefore impacts directly on the health and lifespan of almost 1 billion people, as well as their families. Any such challenge must have a rigorous scientific basis.

In a recent study, Johnson et al. claimed that LCCBs significantly increased the risk of heart failure^7^. This conclusion was based solely on their analysis of hypertensive patients’ medical records, obtained by simply categorizing hypertensive patients into those on LCCBs versus other anti-hypertensive drugs, and then calculating the % in each group that showed heart failure. Mechanistically, they attributed this to the ability of LCCBs such as amlodipine to activate CRAC channels in various non-excitable cells when applied at microMolar concentrations, a conclusion drawn mainly by using the calcium indicator dye fura-2. They also reported that 0.5 μM amlodipine, which barely increased Ca^2+^ signals, interacted synergistically with platelet derived growth factor to increase proliferation of cultured vascular smooth muscle cells. The authors concluded: “Our data indicate caution against the use of LCCBs in elderly patients or patients with advanced hypertension and/or onset of cardiovascular remodelling, where levels of STIM and ORAI are elevated’. This broad and vague statement (vague because advanced hypertension is not a term used by cardiologists or physicians in general) questioning the use of LCCBs in hypertension was reaffirmed in a Penn State University Press release where Dr Trebak stated: “L-type calcium channel blockers are one of the most widely prescribed drugs to treat hypertension, yet we have found that these drugs may cause the same type of damage they are intended to prevent.”

In contrast to Johnson et al.^7^, several prior studies by epidemiologists showed that LCCBs were associated with reductions in stroke and other major cardiovascular events^8–10^.Furthermore, a substantial body of literature has demonstrated that microMolar concentrations of LCCBs block store-operated Ca^2+^ entry^11–15^, rather than activate the pathway as reported by Johnson et al^7^. LCCBs such as amlodipine have also been found to release Ca^2+^ from the endoplasmic reticulum, subsequently reducing Ca^2+^ release to thapsigargin, and to inhibit cell proliferation in vascular smooth muscle^16–21^ instead of stimulating it^7^.

A further area of concern comes from the use of the fluorescent dye fura-2 to measure cytosolic Ca^2+^ in studies with amlodipine. Other groups have reported complex effects of amlodipine on fura-2-based measurements. Asai et al. (2008) found that amlodipine inhibited cytosolic Ca^2+^ signals measured with fura-2 in a concentration dependent manner, with near complete suppression at 10 μM^22^. Because this effect was not mimicked by other LCCBs, the authors concluded “When the effect of amlodipine on intracellular Ca^2+^ concentration is assessed, fura-2 fluorospectrometry should not be used due to fluorescent interaction between amlodipine and fura-2.”^22^ Quentin et al. found substantial intracellular accumulation of amlodipine in organelles, particularly lysosomes, and this led to enhanced emission following excitation at wavelengths used for fura-2, raising further concern with the use of fura-2 with amlodipine^23^. Johnson et al. acknowledged the problems associated with combining fura-2 measurements with amlodipine^7^ but nevertheless used the combination in 23 out of 24 experimental panels when they measured cytosolic Ca^2+^ in their paper. In one set of experiments, they measured cytosolic Ca^2+^ with fluo 4 and reported Ca^2+^ signals to microMolar concentrations of amlodipine, but they did not show this was due to Ca^2+^ entry^7^.

Our recent study^24^ is in good agreement with those earlier works that reported fura-2 was unsuitable for measuring cytosolic Ca^2+^ in the presence of amlodipine. We found that amlodipine was autofluorescent in extracellular solution but became several fold-more fluorescent when it rapidly accumulated within intracellular compartments within seconds to minutes of exposure^24^. Amlodipine washed out of cells very slowly, over tens of minutes, meaning the intracellular signal emanating from its compartmentalisation could not be removed during the time course of a typical cytosolic Ca^2+^ experiment. At the excitation wavelengths for fura-2 (340 and 380 nm), the emission from intracellular amlodipine was considerably higher than for Ca^2+^-fura-2, suppressing the latter signal. Using a longer excitation wavelength dye (Cal 520), we failed to see any Ca^2+^ influx to concentrations of amlodipine of 1 μM or less^24^. At higher concentrations, consistent with many other studies, we observed polypharmacological effects including Ca^2+^ release from thapsigargin-sensitive Ca^2+^ stores and inhibition of store-operated Ca^2+^ entry.

In collaboration with cardiac epidemiologists, our carefully controlled meta-analysis of published clinical trials and a prospective real-world analysis of patients prescribed single antihypertensive agents both showed that dihydropyridines were not associated with increased heart failure or other cardiovascular disorders^24^. By contrast, several well-established confounders of heart failure were not considered in the analysis of patients’ medical records by Johnson et al, raising concerns with their interpretation of the data^7^.

Therapeutic concentrations of free amlodipine in the plasma of patients is in the sub nM to low nM range and increases to tens of nM are associated with toxicity, coma and death^25,26^. The microMolar concentrations that appear to open store-operated Ca^2+^ channels therefore are of questionable clinical relevance.

Trebak et al.^27^ have responded to our findings and raise points to support their earlier conclusions. We do not find their arguments convincing. To address these points, we have carried out several additional experiments which we believe bring clarity to the issues being discussed. These data combined with a detailed analysis of data in Johnson et al.^7^ and Liu et al.^28^, identify several misconceptions and flaws in the arguments by Trebak et al^27^, as well as demonstrably wrong calculations that combine to cast serious doubts on all their main conclusions.

## Methods and Materials

### Cell Culture

HEK293 cells (ATCC, CRL 1573) and RBL2H3 cells (ATCC, CRL 2256) were cultured in Dulbecco’s minimum essential medium (DMEM) supplemented with 10% heat-inactivated fetal bovine serum and 2 mM glutamine and maintained in a humidified 95% air, 5% CO_2_ incubator at 37°C, as described^24^. In general, in preparation for cytosolic Ca^2+^ measurements, fluorescence imaging or confocal microscopy, these cell types were subcultured onto 30 mm round glass coverslips (#1.5 thickness) and maintained in culture for an additional 36-48 h. Experiments with HEK293 cells, were performed with a HEPES-buffered salt solution (HBSS: NaCl 120; KCl 5.4; MgCl_2_ 0.8; HEPES 20; CaCl2 1.8 and glucose 10 mM, with pH 7.4 adjusted by NaOH). For RBL2H3 cells, a slightly modified HEPES-buffered salt solution was used (HBSS: NaCl 145; KCl 2.8; MgCl_2_ 2; HEPES 10; CaCl_2_ 2 and glucose 10 mM, with pH 7.4 adjusted by NaOH).

### Cell Transfection

HEK293 cells were plated in a 6-well plate on Day1. On Day 2, cells were transfected using Lipofectamine 2000 (Invitrogen; 2 μl per well) with cDNA (0.5 μg/well) for EYFP tagged STIM1 and CFP tagged Orai1. After a 6 hr incubation period, the medium bathing the cells was replaced with complete DMEM and maintained in culture. On Day 3, cDNA treated cells were transferred to 30 mm glass coverslips in preparation for single cell Ca^2+^ measurements (as described below), which were performed on Day 4 or 5. At the start of each Ca^2+^ measurement, cells overexpressing EYFP-STIM1 and CFP-Orai1 were identified by the fluorescence from EYFP-Stim1.

### Isolation of Human peripheral blood mononuclear cells (PBMCs)

Blood samples were collected from ‘Control Subjects’ and ‘Test Subjects’ at the Clinical Research Unit (NIEHS) with full consent. To isolate PBMCs, whole blood samples were collected into a BD Vacutainer® CPT™ Mononuclear Cell Preparation Tube containing Sodium Heparin (BD Bioscience, Catalog No:362753). The samples were centrifuged at 1800 x g for 5 min, resulting in mononuclear cells being located in between plasma and the polyester gel. The layer of mononuclear cells was transfered to a 50ml falcon tube, and the cells washed twice with PBS. PMMCs were then resuspended in HBSS containing 2mM CaCl_2_ (HBSS:145 mM NaCl, 2.8 mM KCl, 2 mM MgCl_2_, 10 mM D-glucose, 10 mM HEPES, 0.1 mM EGTA, 2mM CaCl_2_ pH 7.4) and quickly plated on 35 mm glass bottom dish (MATTEK, Part No: P35G-1.5-14-C).

The MATTEK dish was then placed on the stage of a Zeiss AxioObserver epifluorescence microscope (Carl Zeiss Inc, Oberkochen, Germany) equipped with a Zeiss Plan-Apochromat objective (20x/0.8 NA) and a Colibri 7 LED light source. To detect and visualize amlodipine fluorescence, the 385nm LED was used for excitation with a 370-400 nm excitation filter coupled with an emission filter of 500-530nm. A Hamamatsu ORCA Flash C11440 camera was used to collect fluorescence images with binning set to 2×2 and a 50ms exposure. Additionally, time series images we acquired with Definite Focus engaged to eliminate focal drift. Images are taken every 1 min. The time between the onset of centrifugation and the start of the imaging was < 8 minutes.

### Single Cell Calcium Measurements with UV ratiometric calcium indicator

Fluorescence measurements were made with HEK293 cells loaded with the Ca^2+^-sensitive dye, fura-5F, as described previously described^29^. Briefly, cells plated on 30 mm round coverslips and mounted in a Teflon chamber were incubated in DMEM with 1 μM acetoxymethyl ester of fura-5F (Fura-5F/AM, Setareh Biotech, Eugene, OR) at 37° C in the dark for 25 min. Cells were bathed in HEPES-buffered salt solution (HBSS in mM: NaCl 120; KCl 5.4; Mg_2_SO_4_ 0.8; HEPES 20; CaCl_2_ 1.0 and glucose 10 mM, with pH 7.4 adjusted by NaOH) at room temperature. Nominally Ca^2+^-free solutions were HBSS with no added CaCl_2_. Fluorescence images of the cells were recorded and analysed with a digital fluorescence imaging system (InCyt Im2, Intracellular Imaging Inc., Cincinnati, OH). Fura-5F fluorescence was monitored by alternatively exciting the dye at 340 nm and 380 nm and collecting the emission wavelength at 520 nm. Changes in cytosolic Ca^2+^ are expressed as the ‘Ratio (F340/F380)’ of fura-5F fluorescence due to excitation at 340 nm and 380 nm (F340/F380). Before starting the experiment, regions of interests identifying cells were created and 25 to 35 cells were monitored per experiment. For experiments with cells overexpressing EYFP-STIM1 and CFP-Orai1, regions of interests were created by imaging the cells and confirming EYFP-Stim1 fluorescence. Ca^2+^ measurements were performed in HBSS supplemented with 1mM CaCl_2_.

### Visualizing Amlodipine fluorescence and cell energy depletion

HEK293 and RBL2H3 cells were plated on 35 mm glass bottom dish (MATTEK, Part No: P35G-1.5-14-C) and maintained in culture for 24 hrs so that, on the day of experiments, cell confluency was 70-80%. To ‘energy deplete’ cells, cells were bathed in glucose-free HBSS containing a Rotenone/Antimycin cocktail (1μM, Agilent Technologies) and 10 mM 2-deoxy-D-glucose for 60 mins at room temperature. After this incubation period, the bathing solution was switched to HBSS and the MATTEK dish placed on the stage of a Zeiss AxioObserver epifluorescence microscope (Carl Zeiss Inc, Oberkochen, Germany) equipped with a Zeiss Plan-Apochromat objective (20x/0.8 NA) and a Colibri 7 LED light source. To detect and visualize amlodipine fluorescence, the 385nm LED was used for excitation with a 370-400nm excitation filter coupled with an emission filter of 500-530nm. A Hamamatsu ORCA Flash C11440 camera was used to collect fluorescence images with binning set to 2×2 and a 50ms exposure. Additionally, time series images we acquired with Definite Focus engaged to eliminate focal drift. Images are taken every 2 min for 30 min.

### FLIPR cytosolic Ca^2+^ Measurements

Cytosolic Ca^2+^ was monitored in Fluo4-loaded and Cal520-loaded HEK 293 cells using a fluorometric imaging plate reader (FLIPR^TETRA^; Molecular Devices, Inc., Sunnyvale, CA)^24^. HEK 293 cells were plated 24 hrs before use on polyLysine-coated 96-well plates at 40,000 cells/well. On a single 96-well plate, cells were then loaded with either the visible wavelength indicator Fluo4/AM (Rows A-D; 4 μM Fluo4/AM) or Cal520 (Rows E-H; 4 μM Cal520/AM) by incubation in complete DMEM for 45 min at 37 °C. After the indicator loading period, cells were washed twice in nominally Ca^2+^-free HBSS and then bathed in HBSS supplemented with 2mM CaCl_2_. The 96-well plate was placed on the FLIPR observation stage and the indicator-loaded cells were excited at 488 nm with emission fluorescence detected by a cooled charge-coupled device camera via 510–570-nm bandpass filter. Experiments were carried out at room temperature. Changes in cytosolic Ca^2+^ are expressed as the “F/F_o_” of Fluo4 or Cal520 fluorescence, whereby the time course of fluorescence intensities (F) were divided by the initial fluorescence intensity recorded at the start of the experiment (F_o_).

### D1ER and Monitoring Ca^2+^_ER_

To directly measure changes in the levels of ER Ca^2+^ content (Ca^2+^_ER_) we used HEK293 cells stably expressing the ER-targeted D1ER cameleon^30^. These cells were generated by transfecting HEK293 cells using Lipofectamine 2000 (Invitrogen; 2 μl per well) with D1ER cDNA (0.5 μg/well; a generous gift from Dr. Amy E. Palmer). After a 6 hr incubation period, the medium bathing the cells was replaced with complete DMEM and maintained in culture. Subsequently, the population of HEK-D1ER expressing cells were routinely sorted and enriched by flow cytometry.

In preparation for Ca^2+^_ER_ measurements, HEK-D1ER cells were cultured plated on 30 mm round coverslips. On the day of the experiment, the coverslips were mounted in a Teflon chamber, the cells bathed in HEPES that was nominally Ca^2+^-free (no added CaCl_2_), and the chamber mounted on the stage of a Zeiss LSM 780 confocal microscope equipped with a 20x objective (N.A. 0.8W). Fluorescence images were collected with a pinhole set at 10 Airy unit (equivalent to a z-slice of 16 μm). Under these image capture settings, the number of cells in the field of view were 150-200. Three-channel images of D1ER was acquired in a time series with the Zeiss Zen Black software (version 2.3 SP1) The three channels were: Donor channel (excitation 458 nm, emission 464-500 nm), FRET channel (excitation 458 nm, emission 526-597 nm) and a Ratio image that was generated with the following calculation: ((FRET + 1)/(Donor + 1)) * 10. After experiments were completed, the three-channel image was then opened in FIJI (v1.54f) and a macro used to automate the analysis of the time series images. Specifically, the macro would: (i) open the image, (ii) select the donor channel and run the smooth command twice, (iii) after the smooth command, a threshold of the donor channel was used to create a mask of the time series of areas above 25 counts, (iv) a ‘Boolean AND’ command was used with this mask and the Ratio channel to create an new image where the background (non-cell area) now has a pixel value of 0, (v) finally, a threshold was set to this new image to measure the average intensity of pixels greater than 0 thus yielding the mean intensity of only the D1ER expressing cells in the Ratio image. Subsequently, data from the Ratio image was expressed as the “F/Fo” of D1ER signal, whereby the time course of the Ca^2+^ ratio signal (F) were divided by the initial Ca^2+^ ratio signal recorded at the start of the experiment (Fo).

### Measuring NMR characteristics of amlodipine besylate in the absence and presence of Fetal Bovine Serum (FBS)

NMR samples were prepared by diluting FBS and Amlodipine Besylate into phosphate buffered saline (PBS) with 10% 2H_2_O, and 50 mM Sodium trimethylsilylpropanesulfonate (DSS). NMR data was acquired on a 600 MHz Agilent DD2 spectrometer equipped with a cryogenically cooled probe using the Agilent ‘tnnoesy’ pulse sequence with 1s acquisition time, 1s recycle time and 100 ms mixing time. Peak heights of Amlodipine and Besylate were fit with VNMRSYS (Aglient) and Chenomx (Alberta, Canada) software programs to measure concentrations. The spectral characteristics of besylate were not affected by the presence of FBS and provided a convenient internal control to calibrate the change in concentration of amlodipine as the % content of FBS was increased. NMR spectra were therefore scaled to the besylate peak height.

### Measuring spectral characteristics of amlodipine besylate and the fluorescent calcium indicator fura-2

HEK 293 cells were plated 48hrs before use on polyLysine-coated 12-well plates. Cells were cultured in only 8 wells, leaving the remaining 4 wells available for monitoring the spectral properties of bathing solution. Of the 8 wells containing cells, only 4 wells of cells were incubated in DMEM with 4 μM fura-2/AM (fura-2/AM, Molecular Probes, USA) at 37° C in the dark for 45 min. All other wells just contained DMEM. In preparation for the spectral scan, the cells were bathed at room temperature in HEPES-buffered salt solution (HBSS in mM: NaCl 120; KCl 5.4; Mg_2_SO_4_ 0.8; HEPES 20; CaCl_2_ 10 and glucose 10, with pH 7.4 adjusted by NaOH) and then treated with one of four different conditions: (1) DMSO, (2) 2μM TG, (3) 20 μM amlodipine (4) 2 μM TG + 20 μM amlodipine. After a 15 minute incubation period, fluorescence excitation spectra scan (Excitation 330-400 nm, Emission 510 nm) was recorded and analyzed using a VANTAstar fluorometer (BMG Labtech, Cary, NC, USA). For each of the four conditions described above, spectral data was recorded from wells with ‘fura-2-loaded cells’, ‘cells only, no fura-2’ and ‘solutions’.

### Materials

Thapsigargin was purchased from Alexis (San Diego, CA), fura-5F/AM Setareh Biotech (Eugene, OR), Fluo4/AM and Fura-2/AM (Invitrogen, USA), Cal520/AM (ATT Bioquest, CA), heat inactivated Fetal Bovine Serum (Gibco FBS; Thermo Fisher, USA). Amlodipine Besylate and Diltiazem from Selleck Chemicals (Houston, TX), 30mm round #1.5 coverslips from Bioptechs (Butler, PA), BioCoat Poly-D-Lysine 96-well Black/Clear Flat Bottom TC-treated Microplates (Corning, USA).

### Statistics

Data are presented as mean ± SEM, and statistical evaluation on raw data was carried out using ANOVA and Tukey’s multiple comparisons test (GraphPad Prism).

## RESULTS and DISCUSSION

### 1. Shortcomings in cell proliferation experiments

In their letter, Trebak et al.^27^ point out that we had not repeated their ‘key biological observation’ showing 0.5 μM amlodipine, apparently working through store-operated Ca^2+^ channels, and 0.5 ng/ml PDGF interacted synergistically to increase cell proliferation. We do not believe this was a critical experiment for their two major conclusions^7^, namely that LCCBs i) increased heart failure and ii) activated CRAC channels. Regardless, this “key” experiment in Johnson et al. is fundamentally flawed, raising serious doubts over the authors’ interpretation.

First, Trebak et al.^27^ stress binding of amlodipine to plasma proteins, which is 98%^31^, reducing the drug’s free concentration considerably. By their same logic, amlodipine should bind to proteins in the fetal bovine serum (FBS) they used in their experiments. However, they did not consider this at all. In the cell proliferation experiments, Johnson et al. used 0.5 μM amlodipine and 0.4% FBS^7^. We assessed the extent of amlodipine binding to FBS by using NMR spectroscopy. 0.5 μM amlodipine was too low to provide reliable spectra. Nevertheless, because the amlodipine: FBS interaction follows the principles of mass action, what matters are the relative proportions of each agent. Prominent spectra were seen with 100 μM amlodipine besylate, with clear and well-separated peaks for besylate and amlodipine (Figure 1A). Addition of increasing % of FBS had no effect on the besylate signal but substantially reduced the amlodipine spectra (Figure 1A). We quantified the extent of amlodipine binding by integrating the area under the amlodipine spectra and comparing it with the corresponding and constant besylate signal. The Table in Figure 1B shows how this ratio changed with alterations in the amlodipine:FBS ratio, and how this impacted the free amlodipine concentration. As the amlodipine: FBS ratio decreased, free amlodipine concentration also declined (Figure 1B). Because 0.5 μM amlodipine, the concentration used by Johnson et al in Figure 1K of their study, failed to give reliable spectra, we estimated the free amlodipine concentration using extrapolation. A plot of free amlodipine concentration against amlodipine:FBS ratio (Figure 1C) revealed an estimated free amlodipine concentration of ∼1% of total amlodipine with a ratio of 1.25:1 (equivalent to the 0.5 μM amlodipine:0.4%FBS in Johnson et al.^7^) Therefore the free amlodipine concentration used in Johnson et al.^7^ would not have been 0.5 μM but likely to be ∼5 nM. However, there may be some non-linearity in the extrapolations, and so the concentration indicated above is an estimate. Nevertheless, even if one attributes an extremely improbable situation wherein no further amlodipine binds to FBS beyond a ratio of 50:1, the amlodipine concentration in Figure 1K of Johnson et al.^7^ would still only be 165 nM. This is important because 0.5 μM amlodipine was the lowest concentration that gave a Ca^2+^ signal^7^, although this signal was miniscule (see below). Therefore, the free amlodipine concentration in the proliferation experiments of Johnson et al. would have well below the threshold for activating CRAC channels^7^. This is not therefore compatible with their interpretation that 0.5 μM amlodipine activated CRAC channels and this synergised with 0.5 ng/ml PDGF to increase cell proliferation in FBS.

**Figure 1.**
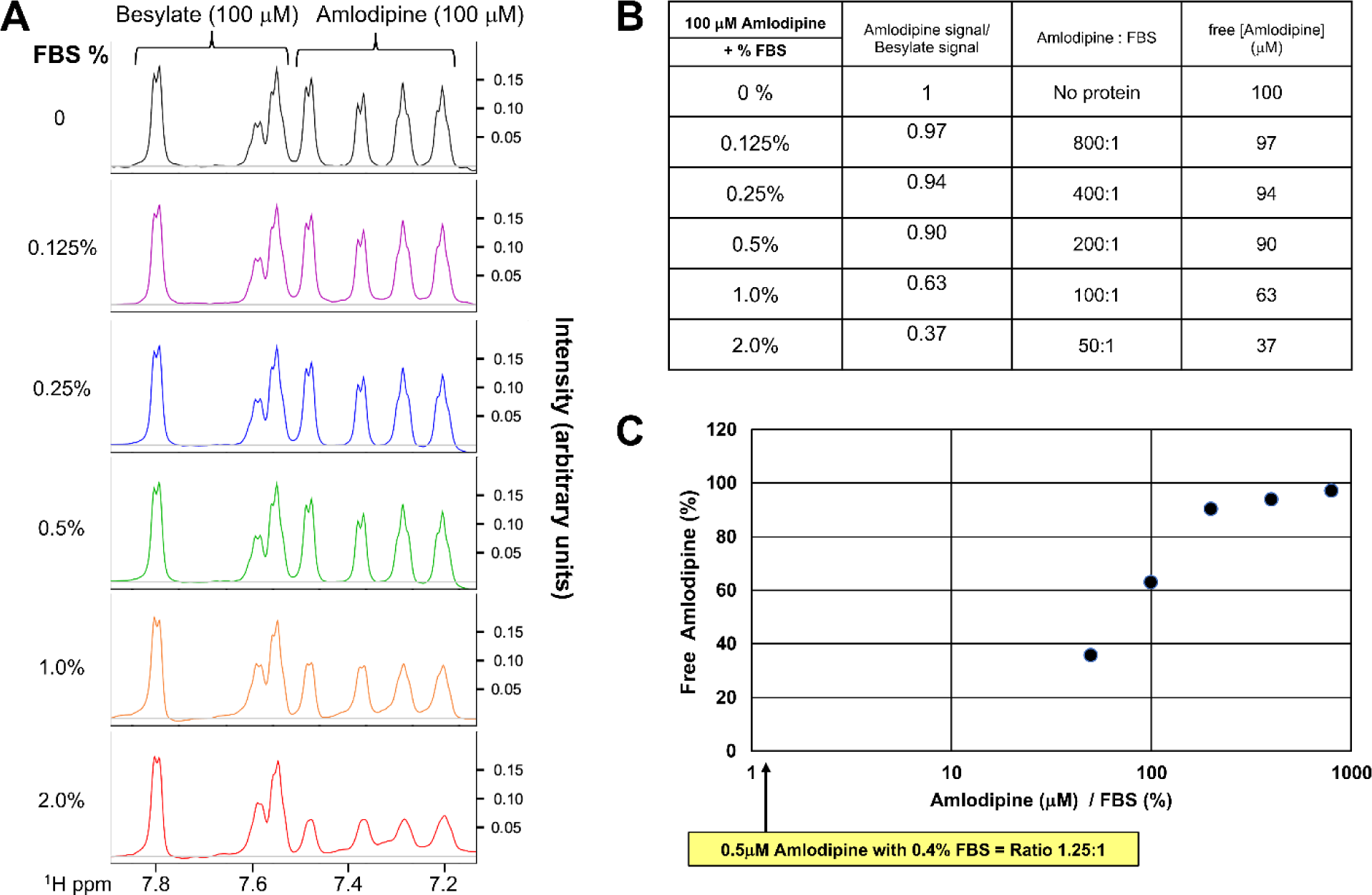
Free amlodipine concentration is reduced by FBS. A, Peak analysis of amlodipine (100μM) and besylate (100μM) in the presence of different % FBS. The spectral characteristics of besylate were not affected by the presence of FBS and provided a convenient internal control to calibrate the change in concentration of amlodipine, as the % content of FBS was increased. B, Table shows how the amlodipine:besylate signal (derived from the peak analysis) changes as a function of % FBS. The corresponding free amlodipine concentration is shown in the right-hand column. The input amlodipine concentration was 100 μM throughout. These data were derived from three independent experiments. C, The graph plots free amlodipine (% of total amlodipine) versus amlodipine:FBS ratio. Based on this plot, estimating the free amlodipine concentration under conditions of 0.5 μM amlodipine mixed with 0.4% FBS (an amlodipine:FBS ratio of 1.25:1) would suggest a free amlodipine of about ∼1%, or 5 nM.

Second, synergy is generally defined as an interaction between two or more components in a system that gives rise to a response that is greater than the sum of each component. Trebak et al.^27^ refer to Figure 1K in Johnson et al.^7^ as key evidence of synergy between 0.5 μM amlodipine and 0.5 ng/ml PDGF, but they provide no test of this. The sum of 0.5 μM amlodipine alone and 0.5 ng/ml PDGF alone looks very similar to their combination (Figure 1K). We therefore accessed the data in the Supplemental Information of Johnson et al. ^7^, related to Figure 1K. We tested for synergy using a 2-way ANOVA test, with the null hypothesis being no difference between synergy and additivity. As shown in the Table 1 below, we found that the results in Figure 1K, using the raw data provided by Johnson et al^7^ can be entirely explained by additivity between 0.5 μM amlodipine and 0.5 ng/ml PDGF.

**Table 1.** Statistical analysis of ‘Interaction’: Synergism vs Additivity. The Table compares the statistical significance between synergy and additivity, using data provided in the Dataset from ref. 7 (<u>pnas.2007598117.sd02.xlsx</u>), related to Figurer 1K in that paper. The analysis shows the data are explained by additivity not synergy.

| Interaction: Synergism vs. Additivity | P value |
| --- | --- |
| 24 <u>hrs</u> | 0.6559 |
| 48 <u>hrs</u> | 0.2997 |
| 72 <u>hrs</u> | 0.4462 |

We also note that only one concentration of one LCCB was used in these proliferation experiments^7^ and no evidence was provided to show the effect of amlodipine indeed reflected Ca^2+^ entry through CRAC channels.

### 2. Contradictory findings between the authors’ papers, leading to questionable clinical relevance

The total concentration of amlodipine in the plasma of patients is in the range of 10-50 nM. With ∼98% protein binding, the *free concentration* of amlodipine is 0.1-0.5 nM and it is the free concentration that determines the volume of distribution of a drug.

In their 2019 Oncogene paper, Trebak and colleagues used ML-9, an inhibitor of STIM aggregation^28^. They stated that ML-9 “inhibit SOCE through a store-independent mechanism by causing rapid reversal of STIM1 aggregation.”. They then reported that ‘50 μM ML-9 was unable to inhibit amlodipine-activated Ca^2+^ entry (Figure 2F in ref. ^28^), further supporting that the effect of amlodipine on Orai1 channels is independent of store depletion and STIM1 aggregation.’

**Figure 2.**
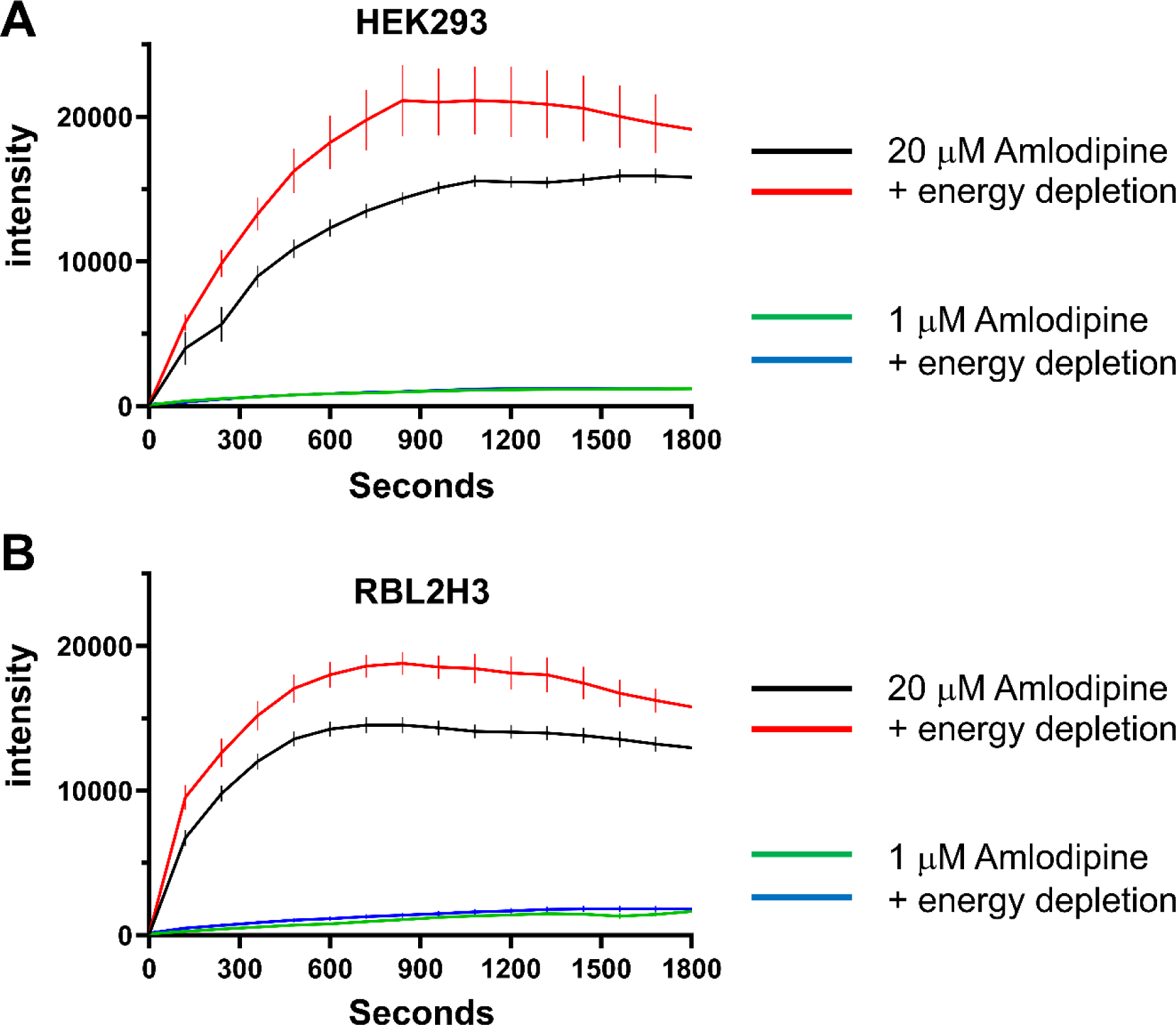
Intracellular accumulation of amlodipine is not an active process. A, Intracellular accumulation of amlodipine was measured over time under energy replete conditions or after energy depletion (see Methods) in HEK293 cells. Amlodipine was applied at either 1 μM (just above the threshold for detecting intracellular accumulation) or at 20 μM. B, As in panel A but RBL2H3 cells were used instead. In both cell types, loading was slightly faster and a little more extensive in energy-depleted cells, suggesting active transport might be required for removal of amlodipine. All data are mean ± SEM for n=5 experiments.

This led to their model (Figure 7 of the Liu et al. 2019^28^), which clearly proposed that amlodipine activated CRAC channels directly, from an external site. Indeed, in the paper, Liu and colleagues stated: “Notably, our results indicated that concentrations of amlodipine to activate Orai1 channels and inhibit cell survival are around 10 μM, which is much higher than the maximum steady state plasma concentration (∼0.04 μM) in clinical studies for its original usage as a hypertension drug.”

We agree with this statement; it is consistent with a vast body of literature that has measured amlodipine concentration in the blood of hypertensive patients. All studies show that therapeutic concentrations are in the sub nM-low nM range, due to protein binding.

In the 2020 Johnson et al, study^7^, the authors reported conflicting results, namely that STIM aggregation was now required for amlodipine activation of CRAC channels. These contradictory findings have not been explained.

The concentrations of amlodipine and other CCBs in the Johnson et al study that apparently activated CRAC channels were in the μM range^7^, with clear responses reported at 20 μM, more than 10^4^ fold greater that therapeutic levels in the blood. The Johnson et al. paper^7^ therefore requires amlodipine (and all other CCBs) to accumulate intracellularly, presumably in the ER, to reach concentrations at least 10,000-fold above the therapeutic levels, for their findings to be clinically relevant. The letter by Trebak and colleagues^27^ create a scenario using pharmacokinetic properties such as volume of distribution and half-life in which such extensive accumulation could be possible. However, the authors provide no evidence in support of this. By contrast, several arguments show that accumulation to the extent the authors propose is debatable.

First, accumulation to this extent could require active transport. Because amlodipine is fluorescent, its accumulation inside cells can easily be monitored^23,24^. Metabolic poisoning of cells failed to affect the rate and extent of intracellular accumulation of amlodipine (20 μM; Figure 2), ruling out active transport. In fact, accumulation was slightly higher after metabolic poisoning, suggesting that active transport might remove amlodipine from intracellular compartments.

Second, different CCBs have vastly different volumes of distribution (Figure 3). There is more than a 25-fold difference in volume of distribution between amlodipine and nifedipine, yet both drugs activate CRAC channels with similar kinetics^7^ and both drugs are effective clinically. By focusing on amlodipine, Trebak et al.^27^ overlook the wide variation in volume of distribution for different LCCBs.

**Figure 3.**
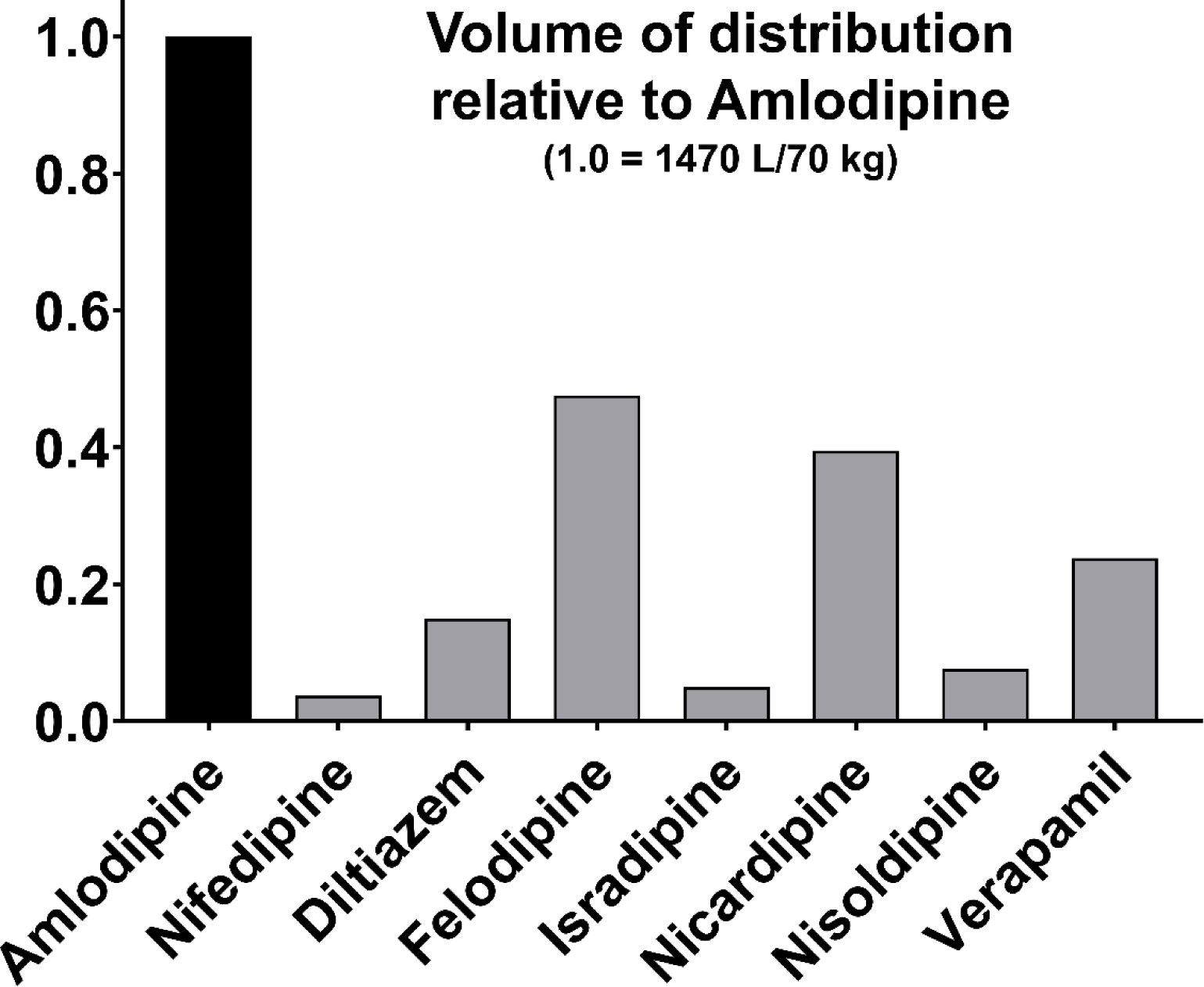
Comparison of volume of distribution of different LCCBs.

Third, a direct prediction of the claim that long-term exposure to amlodipine would lead to significant intracellular accumulation to μM levels, as now proposed by Trebak and colleagues^27^, can be easily addressed using the intracellular fluorescence of amlodipine. We have tested this directly. We drew blood from individuals who had been taking amlodipine continuously for > 15 years for chronic hypertension. We isolated PBMCs and took precautions to minimize cells being exposed to amlodipine-free solution. After centrifugation for 5 minutes, cells were immediately placed under the objective of an epifluorescence microscope and imaged within 3 minutes. The half-time for washout of amlodipine is >45 minutes, and so the PBMCs would have maintained almost all intracellular amlodipine. We failed to see any intracellular fluorescence of amlodipine in the cells, ruling out a possible accumulation of several log orders above plasma levels. However, when we applied 1 or 20 μM amlodipine to the PBMCs from patients or from healthy controls (Figure 4), intracellular accumulation of amlodipine was immediately apparent.

**Figure 4.**
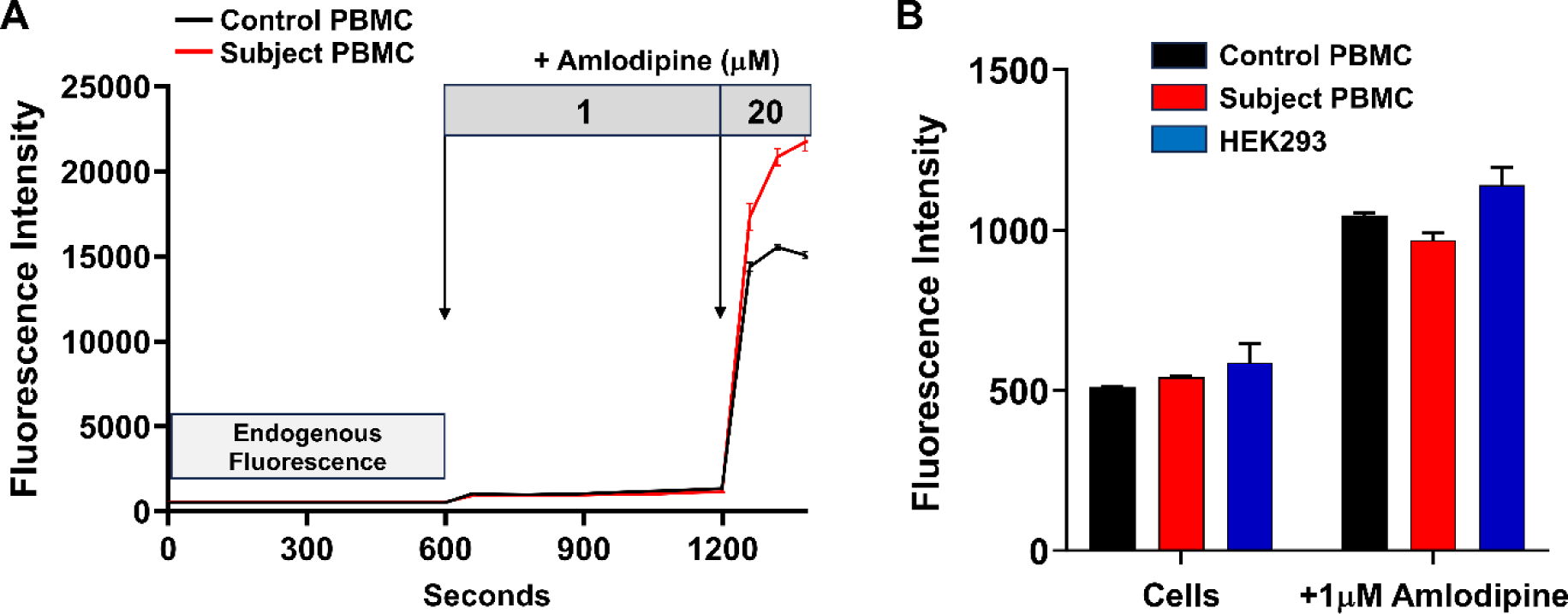
Amlodipine does not accumulate to μM levels inside cells of hypertensive patients. A, Fluorescence of human peripheral blood mononuclear cells (PBMCs), taken from chronic hypertensive and control subjects, was measured (endogenous fluorescence). Then 1μM or 20 μM amlodipine was sequentially applied to confirm whether the drug could indeed accumulate into primary human cells. In chronic hypertensive patients, who had been taking 5 mg amlodipine tablets daily for > 15 years, no endogenous fluorescence was detectable above background autofluorescence (see panel B), but clear increases in fluorescence were observed when 1 μM and 20 μM amlodipine was subsequently applied. B, Aggregate data from several experiments are compared. Cells denotes ‘endogenous fluorescence’ (autofluorescence). +1 μM amlodipine shows the steady state fluorescence levels measured at 900 secs after application of exogenous amlodipine. Endogenous fluorescence in HEK293 cells were measured in parallel experiments under the same experimental condition. HEK293 were exposed to 1 μM amlodipine for 10 minutes and a similar increase in intracellular fluorescence occurred as was seen in subject PBMCs. Note that the HEK293 cells had not been exposed to amlodipine prior to application of 1 μM, unlike the subject PBMCs. Nevertheless, both cell types exhibited similar ‘endogenous fluorescence’, and accumulated amlodipine to similar extents. All data are mean ± SEM for n=5 experiments.

We have therefore failed to find any evidence to support the speculation by Trebak and colleagues^27^ that amlodipine accumulates within cells to levels that are orders of magnitude above therapeutic plasma levels in hypertensive patients. We are further analysing patients to see if there are changes in STIM and Orai, as reported in rodent models.

### 3. Analytical and interpretative issues in observational data analysis

The key conclusion from the Johnson et al. study was that CCBs significantly increased the risk of heart failure compared with other anti-hypertensive agents^7^, and it was this result that led the authors to question the use of CCBs in hypertension. However, the simplistic analyses compounded by errors in group size estimation^7^ limits interpretability. Furthermore, limitations of observational studies are well recognized, including bias and confounding, variable quality and completeness of data and heterogeneity, and because of this, results from observational studies are very low in the evidence hierarchy. We now list some of the analytical errors in the paper.

Their major conclusion that LCCBs increased heart failure compared with all other anti-hypertensive agents was based on their key data that was summarised in Figure 7S and 7T of Johnson et al^7^. The % of patients on LCCBs who developed heart failure was 23.632%, compared with controls who had hypertension but did not develop heart failure. By contrast, patients who developed heart failure on other anti-hypertensive agents (pooled) was 18.548%, a difference of 5.084%. These crucial data (Figure 7S) which was the basis of their key conclusion are reproduced below^7^:

|  | Cases |  | Controls |  |
| --- | --- | --- | --- | --- |
|  | N | % | N | % |
| LCCBs | 10,670 | 23.632 | 45,150 | 76.368 |
| Diuretics | 8,600 | 21.603 | 39,810 | 78.397 |
| β-blockers | 8,450 | 20.575 | 41,070 | 79.425 |
| ACE inhibitors | 5,970 | 15.355 | 38,880 | 84.645 |
| ARBs | 2,140 | 13.342 | 16,040 | 86.658 |
| α-blockers | 2,180 | 18.809 | 11,590 | 81.191 |
| Pooled non-LCCBs | 27,340 | 18.549 | 147,390 | 81.451 |

However, the authors have calculated the % for each group incorrectly, based on a fundamental misunderstanding of percentages. For LCCBs, the % they calculated was Case/Control which is 10,670/45,150, giving 23.632%. This is wrong.

The % should be Cases/(Cases + Controls), which is 10,670/10,670 + 45,150), giving **19.115%.**

Every single entry in Figure 7S of Johnson et al.^7^ is wrong. The corrected table is shown below:

|  | Cases |  | Control |  |
| --- | --- | --- | --- | --- |
|  | N | % | N | % |
| LCCBs | 10,670 | 19.115 | 45,150 | 80.885 |
| Diuretics | 8,600 | 17.765 | 39,810 | 82.2351 |
| β-blockers | 8,450 | 17.064 | 41,070 | 89.936 |
| ACE inhibitors | 5,970 | 13.311 | 38,880 | 86.689 |
| ARBs | 2,140 | 11.771 | 16,040 | 88.229 |
| α-blockers | 2,180 | 15.832 | 11,590 | 84.168 |
| Pooled non-LCCBs | 27,340 | 15.647 | 147,390 | 84.353 |

The authors’ wrong calculations have therefore significantly exaggerated the risk of heart failure between LCCBs and non-LCCBs by as much as 46% (relative risk of heart failure with LCCBs was wrongly stated as an increase of 5.096%, when it should have been 3.468%). affecting the major conclusion of their paper^7^. Figure 7T^7^ is also wrong, as it was derived from the data in Figure 7S. Furthermore, such mathematical mistakes raise major concerns with their analysis of patients’ records in general. These concerns are validated by several other flawed calculations which were important for drawing the conclusions Johnson et al. reached^7^. Because of space limitations, we give only a few examples below. But it is important to note that there are numerous mistakes and wrong calculations.

For example, Figure S10E in Johnson et al.^7^ shows the following Table related to Association of LCCBs with Leiomyosarcomas:

|  | Cases |  | Controls |  |
| --- | --- | --- | --- | --- |
|  | N | % | N | % |
| LCCBs | 490 | 0.567% | 85960 | 99.433% |
| Diuretics | 340 | 0.440% | 76880 | 99.560% |
| β-Blockers | 160 | 0.496% | 32090 | 99.504% |
| ACE Inhibitors | 640 | 0.631% | 100750 | 99.369% |
| ARBs | 560 | 0.541% | 102950 | 99.459% |
| α-Blockers | 200 | 0.618% | 32150 | 99.382% |
| **Pooled non-LCCBs** | 200 | 0.618% | 32150 | 99.382% |

These same data were also analyzed and depicted in the graph in Figure S10F^7^. However, the entry for α-blockers is identical to Pooled non-LCCBs. The Table containing the data used Figure S10F was included in the Supplementary Information by Johnson et al.^7^ and is reproduced below.

| ACE Inhibitors | Diuretics | ARBs | β-Blockers | α-Blockers | **Pooled LCCBs** |
| --- | --- | --- | --- | --- | --- |
| 1.28894424 | 0.897356 | 1.1432716 | 1.0479438 | 0.9163274 | 0.91632736 |
| 1.48069015 | 1.0097502 | 1.3672749 | 1.1834232 | 1.0806682 | 1.0806682 |
| 1.12202898 | 0.7974722 | 0.9559671 | 0.9279742 | 0.7769784 | 0.77697839 |

The entries for α-blockers are given mainly to 7 decimal places, whereas the pooled non-LCCBs are presented to 8 decimal places^7^. Apart from this, the entries are identical. Regardless of how this was brought about, the presented data are incorrect. The graph in Figure S10F^7^ is also wrong, because the entry for α-blockers is identical to that for Pooled non-LCCBs, other than the marker for the latter point being considerably larger.

Another example is Supplementary Figure 10 in Johnson et al.^7^, where there are mistakes in all Tables (S10A, S10C and S10E) and adjacent graphs (S10B, S10D and S10F).

S10A has the following Table:

| **A** | Cases |  | Controls |  |
| --- | --- | --- | --- | --- |
|  | N | % | N | % |
| LCCBs | 1700 | 3.853% | 42420 | 96.147% |
| Diuretics | 1610 | 4.634% | 33130 | 95.366% |
| β-Blockers | 720 | 4.079% | 16930 | 95.921% |
| ACE Inhibitors | 2310 | 4.083% | 54260 | 95.917% |
| ARBs | 1870 | 3.589% | 50230 | 96.411% |
| α-Blockers | 350 | 5.700% | 5790 | 94.300% |
| **Pooled non-LCCBs** | **8560** | **4.051%** | **202760** | **95.949%** |

Pooled non-LCCBs should be the simple addition of all entries **EXCEPT LCCBs.** However, the value of Pooled non-LCCBs for Cases is stated as N= 8560^7^. This is wrong because it is the sum of LCCBs and Pooled non-LCCBs. The correct value is **6860**. Similarly, the Pooled non-LCCBs for Controls is stated as N= 202760^7^. Again, this is wrong. It should be **160340**. The listed % values are incorrect and, because these values directly affect the calculated odds ratio, the graph in S10B is also incorrect^7^.

Similarly, the data in S10C are incorrect.

| **C** | Cases |  | Controls |  |
| --- | --- | --- | --- | --- |
|  | N | % | N | % |
| LCCBs | 610 | 0.706% | 85830 | 99.294% |
| Diuretics | 490 | 0.634% | 76740 | 99.366% |
| β-Blockers | 200 | 0.620% | 32050 | 99.380% |
| ACE Inhibitors | 630 | 0.621% | 100760 | 99.379% |
| ARBs | 610 | 0.589% | 102900 | 99.411% |
| α-Blockers | 240 | 0.742% | 32120 | 99.258% |
| **Pooled non-LCCBs** | **2780** | **0.642%** | **430400** | **99.358%** |

The value of Pooled non-LCCBs for Cases is stated as N= 2780^7^. This is wrong because it is the sum of LCCBs and Pooled non-LCCBs. The correct value is **2170**. Similarly, the Pooled non-LCCBs for Controls is stated as N= 430400. Again, this is wrong. It should be **344570**. The listed % values are incorrect and, so are the calculated odds ratio. The graph in S10D is therefore also wrong^7^.

Another major problem arises in the calculation of heart failure risk between males and females^7^. Using the data provided by Johnson et al. ^7^, we calculated the number and % of heart failure cases compared with controls. These results are shown below:

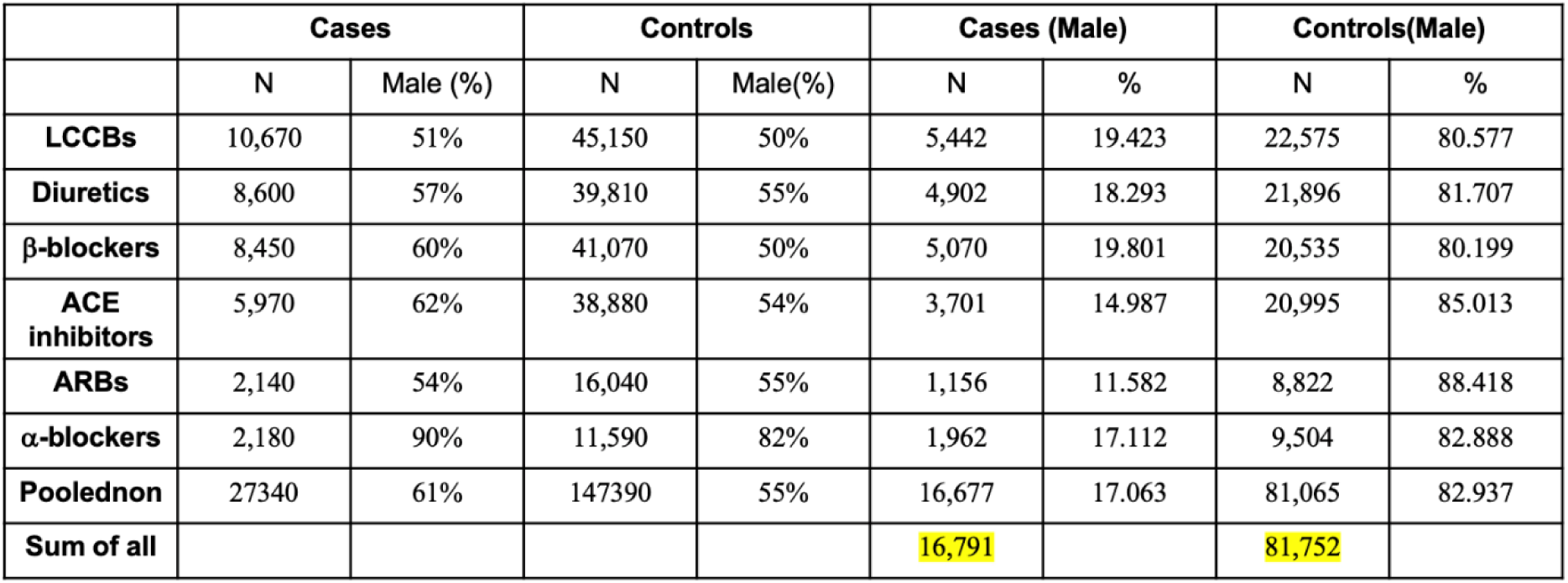

|  | Cases |  | Controls |  | Cases (Male) |  | Controls(Male) |  |
| --- | --- | --- | --- | --- | --- | --- | --- | --- |
|  | N | Male (%) | N | Male(%) | N | % | N | % |
| **LCCBs** | 10,670 | 51% | 45,150 | 50% | 5,442 | 19.423 | 22,575 | 80.577 |
| **Diuretics** | 8,600 | 57% | 39,810 | 55% | 4,902 | 18.293 | 21,896 | 81.707 |
| **β-blockers** | 8,450 | 60% | 41,070 | 50% | 5,070 | 19.801 | 20,535 | 80.199 |
| **ACE inhibitors** | 5,970 | 62% | 38,880 | 54% | 3,701 | 14.987 | 20,995 | 85.013 |
| **ARBs** | 2,140 | 54% | 16,040 | 55% | 1,156 | 11.582 | 8,822 | 88.418 |
| **α-blockers** | 2,180 | 90% | 11,590 | 82% | 1,962 | 17.112 | 9,504 | 82.888 |
| **Poolednon** | 27340 | 61% | 147390 | 55% | 16,677 | 17.063 | 81,065 | 82.937 |
| **Sum of all** |  |  |  |  | 16,791 |  | 81,752 |  |

The N for Pooled Non denotes Pooled non-LCCBs. The N values (cases and controls) depends on whether Johnson et al. calculated the number by summing each entry (highlighted in yellow) or by taking the overall % of males from the total non-pooled number^7^. It is unclear what they did as details are absent from their paper^7^, but the latter approach is incorrect. Using these numbers, and those highlighted in yellow, we calculated the odds ratio for each condition. The data are summarized in the table below:

|  | **OR** | **95%CI** | **z-value** |
| --- | --- | --- | --- |
| **Diuretics** | **1.076684** | **0.989, 1.07** | **1.4681** |
| **β-blockers** | **0.976326** | **0.948, 1.03** | **-0.47798** |
| **ACE inhibitors** | **1.367291** | **1.094, 1.19** | **5.815048** |
| **ARBs** | **1.840206** | **1.217, 1.39** | **7.624565** |
| **α-blockers** | **1.16763** | **1.01, 1.132** | **2.318156** |
| **Poolednon** | **1.17168** | **1.035, 1.10** | **3.970319** |

Our calculations are very different to those reported by Johnson et al. in Supplemental Figure 9D^7^. We find, using the data of Johnson et al., that β-blockers have a higher risk of heart failure in males than LCCBs, whereas diuretics are slightly more protective. This is opposite to what they show in Supplemental Figure 9D^7^. In fact, all our odds ratio calculations are quite different from theirs, using their own data. This is a concern; our analyses have been conducted by Biostatisticians who are experts in this area.

As discussed in Bird et al.^24^, the analysis by Johnson et al.^7^ used a cross-sectional study comparing co-prescription of CCB with heart failure versus heart failure with non-CCB drugs. This is not the preferred method to infer a relationship between any drug and an outcome as this is subject to wide-ranging confounding issues and biases. No differentiation was made between CCBs (verapamil and diltiazem) that have greater cardio-depressant effect and dihydropyridines like amlodipine. Furthermore, there was no adjustment for known confounders such as age, ethnicity, smoking, prevalent cardiovascular disease, obesity, diabetes, or renal disease. The control group in the Johnson et al. study that was not on CCBs might have been on drugs that have a protective effect against heart failure, such as ACE inhibitors, angiotensin receptor blockers or beta-blockers, or combinations thereof. Nor was the duration of treatment taken into account. None of these confounding factors were considered in the Johnson et al. study^7^.

Lack of consideration of confounding factors together with the numerous mistakes in calculations could explain several anomalous findings in Johnson et al^7^.

For example, diuretics are widely prescribed for chronic hypertension and the AllHat study demonstrated that diuretics were more effective than ACE inhibitors and LCCBs in reducing blood pressure and risks of cardiovascular disease^32^. However, analysing the data provided by Johnson et al.^7^ and using the corrected values for heart failure, reveals that diuretics are associated with a highly significant increase in heart failure (p<10e^-10^). Similarly, their corrected data shows that another well-established therapy, namely use of beta blockers, is also associated with a significant increase in heart failure (p<0.0001).

The lack of consideration of confounding issues in the Johnson et al.^7^ study combined with the flaws in their calculations could also explain their unexpected conclusion of sex-based differences in the effectiveness of various anti-hypertensive agents (Supplementary Figure 9D)^7^. This is at odds both with numerous large-scale epidemiological studies and with clinical practice^33,34^.

Trebak et al.^27^ question our prospective real-world analysis of CCBs^24^, arguing that patients were not hypertensive for long enough for vascular remodelling to take place. We fully recognize the limitations of observational studies and we have been very careful in our analysis plan to minimise confounding and bias, as we have described^24^. Our study was conducted on patients with no prior history of cardiovascular disease and who had been newly prescribed one of the 5 classes of antihypertensive drugs as monotherapy^24^, ruling out some of the issues that plagued the Johnson et al. study^7^. Furthermore, the approach adopted by Johnson et al., namely filtering patients’ records simply based on hypertension, would not have considered the duration of hypertension, how long an intervention was administered or whether there were other cardiovascular issues^7^. Their study did not separate patients into those with or without vascular remodelling. Our primary analysis was the systematic review of RCTs (randomized controlled trials) which rank the highest in the evidence hierarchy and we used our observational analyses to explore additional questions that were not possible in the systematic review.

It is important to note that a recent secondary analysis of a randomized clinical trial in adults with hypertension and coronary heart disease risk factors, and which followed up patients for up to 23 years, found cardiovascular disease mortality was similar between groups on diuretics, amlodipine and an ACE inhibitor^35^. There was also no difference between amlodipine and the ACE inhibitor in incidence of heart failure^35^.

Our meta-analyses and prospective real-world study were conducted rigorously following established best practice in the field.

Trebak et al.^27^ point out that some of the data from our rigorous meta-analysis and prospective real-world studies are in agreement with some of their findings, which they seem to take as validation of their approach. We believe any similarity is purely coincidental, in light of their flawed calculations, their comparison of non-comparable groups and lack of consideration of well-established confounders.

### 4. Concerns with amlodipine-induced I_CRAC._

An argument Trebak et al.^27^ make to justify activation of CRAC channels by amlodipine is the patch clamp recording in Figures 2C and 2D of the Johnson et al. study^7^. However, these data are unconvincing. First, the whole cell current is increasing in DMSO prior to administration of amlodipine (Figure 2C^7^). This was not corrected for and leads to an overestimate of the size of the amlodipine-evoked current. Second, the whole cell current induced by amlodipine does not resemble I_CRAC_. I_CRAC_ shows strong inward rectification, with the amplitude of the current increasing steeply at potentials negative to -100 mV^36^. The I-V relationship of the amlodipine-induced current reported by Johnson et al.^7^ looks very different (Figure 5): the I-V relationship is linear to -100 mV (black line) then reaches a plateau beyond -100 mV (yellow line), where the current fails to increase further, despite the increased electrical driving force (Figure 5).

**Figure 5.**
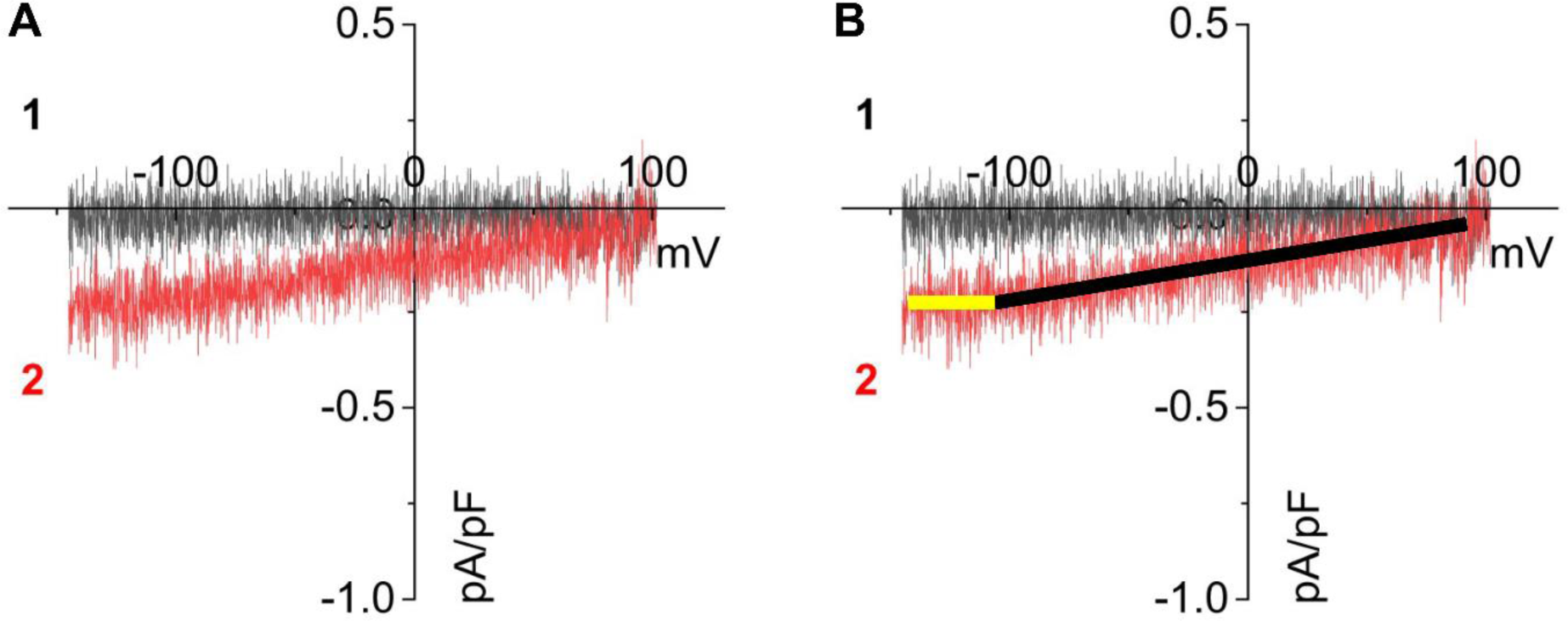
Patch clamp recording of an inward current from Johnson et al^7^ (Figure 2C) is shown. B, the inward current, attributed to I_CRAC_ by Johnson et al, can be fitted by two linear components. Neither component bears any resemblance to I_CRAC_.

Figures 2E and 2F in Johnson et al.^7^ show that 20 μM amlodipine evokes a large monovalent Na^+^ current through CRAC channels in the absence of external divalent cations. However, there are concerns with these data. The kinetics of current activation are quite different in Figure 2E compared with Figure 2C and all cytosolic Ca^2+^ measurements^7^. In the latter experiments, amlodipine-induced Ca^2+^ signals and I_CRAC_ (Figure 2C) started immediately upon exposure, whereas a sizeable delay is seen in Figure 2E. We note Johnson et al. omitted ATP from the patch pipette solution^7^ and ATP is needed to prevent passive store depletion and spontaneous activation of CRAC channels. In divalent-free extracellular solution, CRAC channels exhibit permeability to Cs^+^, with PCs/PNa of ∼0.13^37^. Cs^+^ permeation is widely seen as a clear outward current positive to +60 mV. Johnson et al used a Cs^+^-based pipette solution and one would therefore have expected an outward Cs^+^ current in Figure 2E^7^. This is completely absent from their recording. Finally, the kinetics of activation of the inward current in divalent-free solution is 4-fold faster than in Ca^2+^-containing solution^7^.

Third, the amlodipine-induced currents are small in Ca^2+^-containing solution (mean of -0.2 pA/pF at -100 mV, or -2 to-4 pA whole cell current in a 10-20 pF cell). A small change in seal resistance during application of amlodipine could potentially induce such currents.

Fourth, these data contradict patch clamp recordings from an earlier study by the same group (Zhang et al. 2011^37^). The Zhang et al. study^38^ was cited by Johnson et al.^7^ in the Methods section for how the patch clamp experiments were conducted. In Zhang et al, it is stated in the Methods Section (under Whole-cell patch-clamp electrophysiology, last 3 sentences): “10 μM verapamil was included in bath solution to inhibit L-type calcium channels, and 3 μM nimodipine was added to the bath solution to generally stabilize membrane patches and reach better seals.” ^38^ Johnson et al. report these CCBs activate CRAC channels^7^. If this is true, then the combination of verapamil and nimoldipine should have activated I_CRAC_ in Zhang et al.^38^. However, no such current was observed upon break-in (absence of current after break-in in Figure 1G) nor was a current seen until store depletion a few minutes later^38^. By contrast, Johnson et al, reported CCBs rapidly activated CRAC channels^7^. The two papers show very different effects of CCBs on CRAC channels. Both papers cannot be correct.

It is also important to note that several prior studies have failed to see activation of store-operated Ca^2+^ entry channels to a variety of CCBs in non-excitable cells, including vascular smooth muscle. Some examples are included in Table 2.

**Table 2.** Effects of LCCBs on store-operated Ca^2+^ entry in different cell types. These studies avoided the combined use of amlodipine with fura-2.

| CELL TYPE | LCCB;<br>CONCENTRATION | COMMENTS | REFERENCES |
| --- | --- | --- | --- |
| Vascular Smooth Muscle | Nifedipine; 1 $\mu\text{M}$ | Store-operated $\text{Ca}^{2+}$ current blocked by Nifedipine. Nifedipine did not activate store-operated $\text{Ca}^{2+}$ entry. | 40 |
| Vascular Smooth Muscle | Nifedipine; nM - 100 $\mu\text{M}$ | Nifedipine blocked store-operated $\text{Ca}^{2+}$ entry. $\text{IC}_{50}$ of 24.3 $\mu\text{M}$ Fura-2 | 41 |
| HL-60 | Nifedipine<br>Nicardipine<br>Dihydropyridine analogs | All blocked store-operated $\text{Ca}^{2+}$ entry in $\mu\text{M}$ range<br>Fura-2 | 15 |
| Hepatoma | Nifedipine<br>Nicardipine<br>Verapamil | All blocked store-operated $\text{Ca}^{2+}$ entry.<br>Fluo-3 | 42 |
| Vascular Smooth Muscle | Nifedipine; 1 $\mu\text{M}$<br>Nicardipine; 1 $\mu\text{M}$<br>Nimodipine; 1 $\mu\text{M}$ | 20-30% inhibition of store-operated $\text{Ca}^{2+}$ entry at 1 $\mu\text{M}$ | 43 |
| Astrocytes | Nifedipine; 10 $\mu\text{M}$ | No block of SOCE;<br>Nifedipine did not activate SOCE | 44 |
| Vascular Smooth Muscle | Verapamil +<br>Nifedipine | Neither CCB activated SOCE | 45 |
| U937 | Nifedipine,<br>Nicardipine; 10 $\mu\text{M}$ | Block SOCE;<br>CCBs do not activate SOCE. | 11 |
| Human myometrial cells | Nifedipine; 1-100 $\mu\text{M}$ | No activation of Orai channels | 46 |

### 5. Methodological issues combining fura-2 with amlodipine

Previous studies raised concerns with the use of fura-2 to measure cytosolic Ca^2+^ in the presence of amlodipine^22^, which Johnson et al. acknowledged^7^. Nevertheless, Johnson et al. still used fura-2 with amlodipine in 23 out of 24 cytosolic Ca^2+-^related panels in their paper^7^. Our data^24^, and those of others^22,23^, demonstrate that fura-2 cannot be used to measure cytosolic Ca^2+^ in the presence of amlodipine because amlodipine is i) **highly fluorescent**, ii) is **excited at 340 and 380 nm** wavelengths like fura-2, iii) **accumulates rapidly inside** cells which increases its fluorescence several fold further^23,24^, and iv) yields **emission signals that are several fold** larger than those induced by Ca^2+^-fura-2. Amlodipine therefore dominates the cellular fluorescence signal, masking Ca^2+^-fura-2 responses. This effect is so strong that the Ca^2+^ signal evoked by thapsigargin even in cells overexpressing STIM1 and Orai1 is almost completely suppressed by amlodipine (Figure 6).

**Figure 6.**
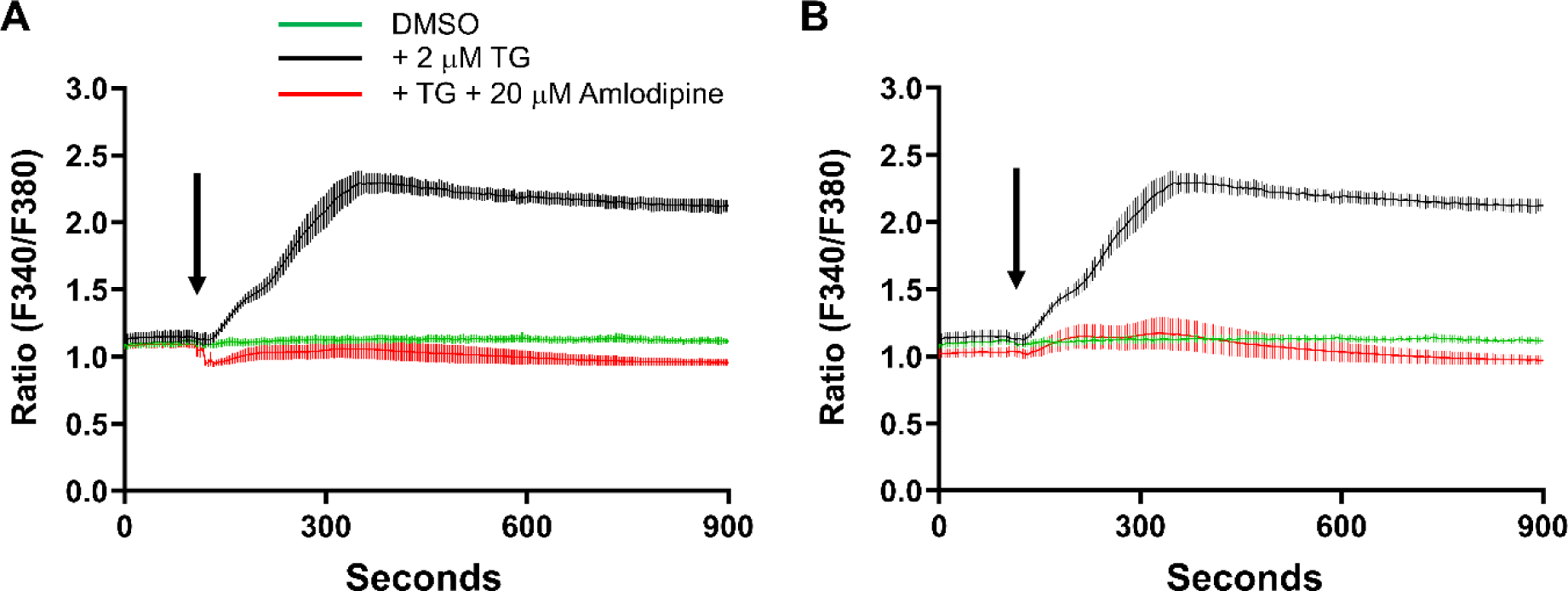
Amlodipine (20 μM) suppresses Ca^2+^ signals to thapsigargin (2 μM) in fura-2-loaded HEK293 cells overexpressing STIM1 and Orai1. STIM1 and Orai1 were overexpressed to increase the size of store-operated Ca^2+^ entry. A, Application of 20 μM amlodipine with 2 μM thapsigargin (red trace) resulted in suppression of the thapsigargin-evoked Ca^2+^ response (black trace). B, Autofluorescence signals measured in extracellular solution at 340 nm and 380 nm (± Amlodipine) were subtracted from the relevant traces in panel A and the new corrected ‘Ratio’ plotted. Despite this ‘correction’, which Trebak et al.^27^ describe as the method they used to eliminate amlodipine fluorescence, the thapsigargin-evoked Ca^2+^ signal remained suppressed. This is because the correction fails to address the major problem of intracellular amlodipine fluorescence. All data are mean ± SEM for n=3 experiments. Note also that amlodipine was co-applied with thapsigargin yet still suppressed Ca^2+^ signals to the latter. This shows that amlodipine accumulates intracellularly sufficiently rapidly to supress Ca^2+^ signals induced at the same time as amlodipine exposure.

We have rigorously obtained excitation spectra for fura-2 and amlodipine, and these are shown in Figure 7. For fura-2-loaded cells (panel A), the blue trace shows the fura-2 spectrum in 10 mM external Ca^2+^. In the presence of thapsigargin (green trace); the 340 signal increases and the 380 decreases, with the isobestic point being at 360 nm, as expected and validating the experimental set-up. The red trace shows the effect of amlodipine in 10 mM Ca^2+^. The signal increases at all wavelengths, including at 340 and 380 nm and at the isobestic point. The amlodipine excitation spectrum overwhelms the fura-2 fluorescence over the entire excitation range. Figure 7B show amlodipine fluorescence in cells not loaded with fura-2 and 7C shows autofluorescence of amlodipine in external solution without cells. Autofluorescence from amlodipine in solution is much smaller than the fluorescence seen in cells, due to the intracellular accumulation of amlodipine, particularly in lysosomes that we and others have described^23,24^.

**Figure 7.**
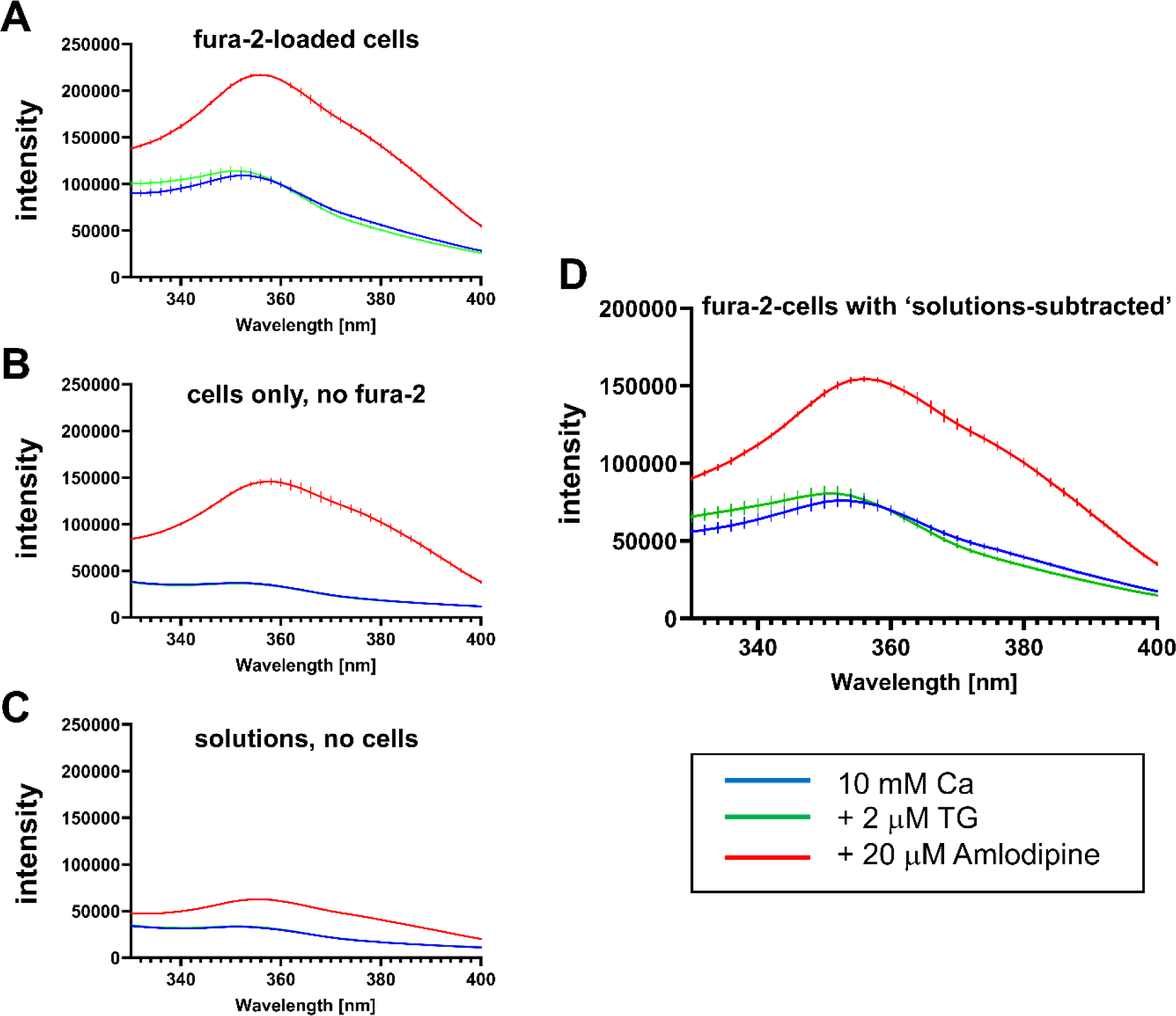
Spectral analysis of fura-2 and amlodipine show that intracellular accumulation of the LCCB overwhelms fura-2 fluorescence at all excitation wavelengths. Excitation spectra (Excitation 330-400 nm, Emission 510 nm) were recorded under four conditions (indicated in the Figure) and from wells that contained: Panel A: ‘fura2-loaded cells’, Panel B: ‘cells only, no fura2’ and Panel C: ‘solutions’. A, Compared to the ‘10mM Ca’ condition (blue trace), treating fura-2-loaded HEK293 cells with thapsigargin increased the fura-2 emission at 340 nm excitation and reduced emission at 380 nm excitation. In the presence of 20 μM amlodipine (red trace), the excitation spectrum measured increased substantially and was enhanced over the entire excitation spectrum occupied by fura-2. B, As in panel A but cells had not been loaded with fura-2. The spectrum to 20 μM amlodipine was only marginally smaller than that in panel A, demonstrating that intracellular accumulation of amlodipine was the dominant source of fluorescence. C, Spectra are compared from extracellular solution alone. D, Subtraction of panel D spectral data from the respective spectra in panel A should eliminate the amlodipine signal, according to the correction protocol applied by Johnson et al^7^, as discussed in Trebak et al^27^. Such correction is wholly inadequate because the amlodipine signal remains and is considerably larger than the fura-2 responses to thapsigargin challenge. All data are mean ± SEM for n=3 experiments.

Trebak et al.^27^ claim that: “Contrary to the assertion of Bird et al., we have recognized the amlodipine fluorescence issue, which we acknowledged, discussed and corrected for (first paragraph in results)4.”

Unfortunately, they did no such thing^27^. The paragraph they refer to simply states: “ and others have noted that LCCBs interfere with the signal of the Ca dye fura-2 and strongly blunt its fluorescence, thus underestimating the magnitude of LCCBs-activated Ca signals (37.38). Therefore, we used fura-2 (being ratiometric) with 10 mM extracellular Ca^2+^ to enhance the driving force and amplify Ca^2+^ signals”.

In the Supplementary Material, under Single Cell Ca^2+^ imaging, Johnson et al.^7^ state: “Addition of amlodipine causes an equal increase of emitted fluorescence upon excitation at 340 and 380 nm^4^, and this autofluorescence from the compound was subtracted.” This is the only information on how they corrected for the amlodipine artefact.

Johnson et al’s solution to this problem was first, to raise external Ca^2+^ to 10 mM and second, to subtract only the amlodipine autofluorescence from the extracellular solution^7^. These measures do not address the problem of intracellular amlodipine fluorescence and are wholly inadequate.

Johnson et al.^7^ have only subtracted the autofluorescence of amlodipine, which is the fluorescence seen in extracellular solution. They were unaware of the more significant problem of rapid intracellular accumulation of amlodipine that dominates the cellular fluorescence, even in the presence of fura-2 and thapsigargin. Figure 7D shows fluorescence spectra in fura-2-loaded cells, after subtraction of the amlodipine autofluorescence in solution (Figure 7C), as was done by Johnson et al.^7^. According to Johnson et al.^7^, subtracting the intrinsic amlodipine autofluorescence should have generated a spectrum that falls within the bounds of the blue and green traces. This is clearly not the case (Figure 7D); the subtraction has failed to remove the more significant signal arising from intracellular accumulation of amlodipine. Because of the rapid accumulation of amlodipine intracellularly (within seconds to minutes), its fluorescence overwhelms the entire fura-2 spectrum and because of its very slow washout from cells (tens of minutes), amlodipine cannot be used in combination with fura-2 to measure Ca^2+^ signals.

Trebak et al.^27^ argue that only the ratio of fura-2 is important, stating that the “individual wavelengths are meaningless.” We fundamentally disagree with this claim. First, the emission ratio (from 340 and 380 nm excitation) is directly derived from the individual wavelengths. Second, emissions at 340 and 380 nm excitation are important diagnostics for any underlying issues, such as fura-2 quenching, that could lead to a misleading or false ratio signal. Best practice requires rigorous examination of these signals to accurately assess the fidelity of recordings. Third, knowing the size of the Ca^2+^-fura-2 signal at each wavelength relative to amlodipine fluorescence at the same excitation wavelengths in each cell is critical. A large amlodipine signal masks the fura responses at each wavelength, making it undetectable. Johnson et al’s^7^ lack of consideration of individual wavelengths has led to artefacts that question the validity of their ratio measurements, as described in the following section.

### Validation experiments by Johnson et al. are hard to interpret

Johnson et al^7^ cite reference 37, their Oncogene paper (Liu et al., 2019^28^), in which they first reported effects of amlodipine on store-operated Ca^2+^ entry, primarily using fura-2. Trebak et al.^27^ refer to this paper^28^ as the one containing the key controls that justify their combined use of fura-2 and amlodipine. The relevant data are Supplementary Figures S2J and S2K^28^. Figure S2J is critical, as it shows emission to the individual 340 and 380 nm excitation wavelengths. As we show below, the data are uninterpretable.

Fura-2 is a ratiometric dye. When cytosolic Ca^2+^ rises, the emission at 340 nm excitation increases whilst the emission at 380 nm decreases, translating into an increase in the 340/380 ratio. For this ratio to be meaningful, it is critical that the emission at 340 and 380 nm excitation are taken as closely together as possible, in order to be recording signals at the same time. A temporal offset of more than a few hundreds of milliseconds renders the ratio meaningless.

Figure S2J from Liu et al.^28^ is reproduced below. We managed to extract the raw data points from the published PDF and the emission signals to 340 and 380 nm excitation wavelengths superimposed with their Figure S2J^28^, showing the data extraction fully mirrors the original published traces. Analysis of the recordings reveal several fundamental problems that raise major concerns with how Liu et al^28^ and Johnson et al.^7^ measured Ca^2+^ with fura-2 and amlodipine. These are shown in Figure 8 and discussed below.

i. Shortly after amlodipine addition (in Ca^2+^-free solution), emission at both 340 and 380 nm excitation increase, but follow very different kinetics. The 340 signal increases in a biphasic manner, with a t_1/2_ of 58.54 seconds. However, the 380 signal increases mono-exponentially with a t_1/2_ of 7.45 seconds, an ∼8-fold temporal mismatch.
ii. When Ca^2+^ is readmitted, the 340 signal increases instantly, followed by a slower secondary rise. The 380 nm signal, which should decrease if cytosolic Ca^2+^ is elevated, **also increases** and remains above prestimulation levels for 71 seconds. It then decreases but increases again and returns to pre-stimulation levels 8.86 minutes after addition of Ca^2+^. This occurs despite the 340 signal remaining elevated. The 340 and 380 signals therefore poorly correlate and do not follow expected cytosolic Ca^2+^ changes.
iii. In 50 μM 2-APB, which blocks CRAC channels, there is a huge temporal mismatch between 340 and 380 signals. The 340 signal decreases with a t_1/2_ of 28.92 seconds, whereas the 380 nm signal rises with a t_1/2_ of 102.66 seconds, almost 4-fold slower.
iv. Most troubling, when the authors wash out amlodipine (Figure 8), the 340 nm signal increases and the 380 signal decreases^28^. As amlodipine increases both 340 and 380 signals when it is applied at the start of Figure S2J; see our Figure 8), one would expect the converse when the drug is removed. The decrease in 380 nm therefore makes sense. But the huge increase in the 340 nm signal is hard to understand. Furthermore, the decrease in the 380 signal has a t_1/2_ of 213.6 seconds, whereas the increase in the 340 signal has a t_1/2_ more than an order of magnitude larger. The opposing changes on washout of amlodipine is troubling, as are their vastly different kinetics.

**Figure 8.**
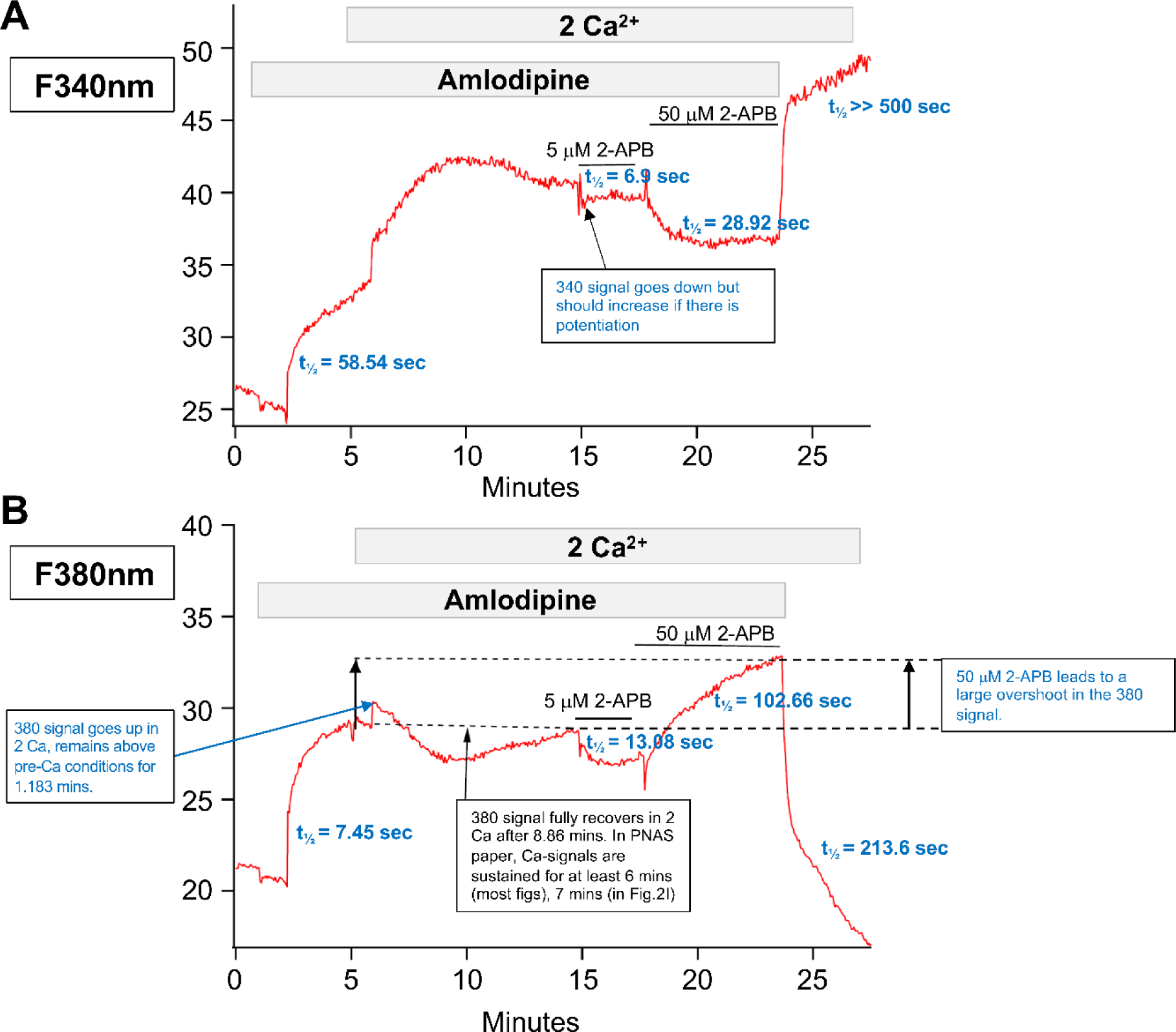
Analysis of the amlodipine-fura-2 validation experiment by Liu et al. reveals major problems in data acquisition and interpretation. A, Analysis of various components of the emission signal following 340 nm excitation. B, As in A but the signal following 380 nm excitation is shown. The kinetics of change at each wavelength differ by several fold. See Figure and text for specific details.

Despite assertions in Johnson et al.^7^, and the perspective by Trebak et al. ^27^, we believe their experiments do not validate the combined use of amlodipine and fura-2 at all. Instead, the marked mismatching between 340 and 380 signals, the fact that the signals do not match expected Ca^2+^ changes, and the large and continuous **increase** in fluorescence at 340 nm upon washout of amlodipine all raise serious concerns with data acquisition, interpretation and whether the authors are indeed able to measure meaningful Ca^2+^ signals in these experiments.

Trebak et al.^27^ seem to argue that, because they did not see the small 340/380 ratio changes induced by amlodipine in STIM KO cells, then their approach was valid. However, these experiments used fura-2 to measure cytosolic Ca^2+^, which, as shown above, is highly problematic. The approach is flawed and using the same approach to show a control does not validate the approach.

#### Data from other dyes and the literature are consistent with our findings

In Bird et al., we showed that Cal520 was suitable for measuring cytosolic Ca^2+^ without contamination by amlodipine fluorescence^24^. We have also used fluo 4. In both cases, we fail to see any Ca^2+^ signals to ≤1 μM amlodipine or other CCBs (Figure 9), even in cells overexpressing STIM1 and Orai1 and where the thapsigargin-evoked responses are increased substantially.

**Figure 9.**
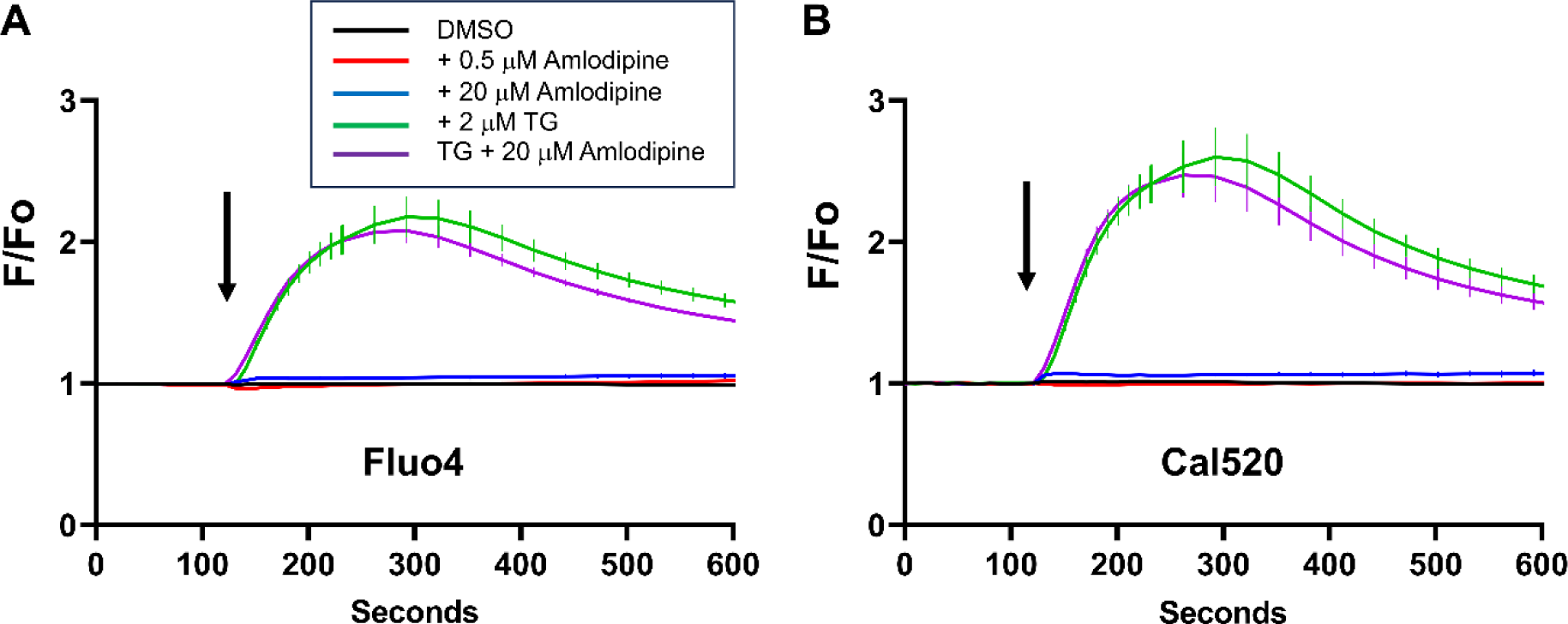
Effect of amlodipine on cytosolic Ca^2+^ in cells expressing longer excitation wavelength dyes, and with spectral properties which do not overlap with amlodipine excitation wavelengths. A, in fluo4-loaded cells, 0.5 μM amlodipine failed to increase cytosolic Ca^2+^. 20 μM amlodipine evoked a miniscule response. Whilst thapsigargin elicited a substantial Ca^2+^ signal, it was reduced in the presence (co-addition) of 20 μM amlodipine at later time points. 20 μM amlodipine inhibits store-operated Ca^2+^ entry modestly, accounting for the reduced Ca^2+^ signal at later times, when the channels have opened. B, as in panel A but Cal520 was used instead. Data in B are in excellent agreement with our earlier results using Cal520^24^. All data are mean ± SEM for n=6 experiments.

Johnson et al. reported Ca^2+^ signals to amlodipine in fluo 4-loaded cells^7^. We do not have an explanation for this discrepancy with our data, but we note several other groups have failed to see activation of SOCE to CCBs using fluorescent dyes or electrophysiology (Table 1), consistent with our results. In Figure 1F, Johnson et al. report a miniscule Ca^2+^ rise to 0.5 μM amlodipine in fluo-4 loaded cells^7^. However, the vehicle control (black trace) which records basal Ca^2+^, is decreasing over time, giving the impression of a larger Ca^2+^ signal to 0.5 μM amlodipine than is the case^7^. We also note that amlodipine was not applied in Ca^2+^-free solution in any of their fluo-4 experiments, which is necessary to rule out intracellular release of Ca^2^, especially as the authors reported masking of Ca^2+^ release by thapsigargin in fura-2-loaded cells in the presence of amlodipine^7^ or other CCBs (data not shown), even in cells overexpressing STIM1 and Orai1 and where the thapsigargin-evoked responses are increased substantially.

#### Polypharmacological effects of CCBs on calcium signaling

Trebak et al.^27^ state that much of our calcium signaling data^24^ are compatible with theirs. This is disingenuous for several reasons: i) Our data show that amlodipine concentrations ≤1 uM do not evoke a detectable Ca^2+^ signal in either fluo-4 or Cal520-loaded cells, which avoid the artifact associated with use of fura-2. ii) At high concentrations (20 μM and above), amlodipine has multiple effects. It releases Ca^2+^ from the thapsigargin-sensitive ER and inhibits store-operated Ca^2+^ entry^24^. There is a small window where amlodipine evokes weak Ca^2+^ influx, but we think this is mainly canonical gating through store depletion^24^ because it is only seen with amlodipine concentrations of ∼20 μM and which associate with Ca^2+^ release from the stores and which do not fully block store-operated channels. (iii) Nifedipine failed to evoke Ca^2+^ influx over a concentration range of 0.5 μM-100 μM^24^, contrasting with Johnson et al who stated that all CCBs activate CRAC channels^7^. These are major differences that lead to different conclusions. As we discuss in our paper^24^, several groups have published data that are entirely consistent with ours including depletion of the thapsigargin-sensitive Ca^2+^ store by amlodipine, inhibition of CRAC channels by CCBs with IC_50_s in the μM range, a failure of CCBs to activate Ca^2+^ entry through CRAC channels (Table 2) and inhibition of cell proliferation by amlodipine, in contrast to Johnson et al. These are all cited in Bird et al. ^24^

Johnson et al.^7^ reported activation of CRAC channels to 20 μM amlodipine and, using a Ca^2+^ probe targeted to the ER, they did not observe any store depletion to this concentration of amlodipine. We measured the extent of store release to amlodipine by assessing the size of a subsequent response to the ER Ca^2+^-mobilising agent ionomycin^24^. Whilst 30 and 40 μM amlodipine caused clear store Ca^2+^ release, 20 μM amlodipine did not evoke a significant change^24^. One possibility is that there is a threshold > 20 μM amlodipine which is required for Ca^2+^ release. Alternatively, 20 μM amlodipine releases Ca^2+^ from the ER but this is modest, so Ca^2+^ extrusion by the Gd^3+^-sensitive PMCA pump prevents cytosolic Ca^2+^ from rising. We therefore carried out the experiment described in Figure 5F and 5G of Bird et al. ^24^, which show significant Ca^2+^ release to 20 μM amlodipine when Ca^2+^ extrusion was reduced^24^.

We acknowledge that Johnson et al. measured ER Ca^2+^ directly using GCaMP6-150^7^, whereas we assessed ER Ca^2+^ content indirectly^24^. We therefore repeated the experiments with a widely used ER Ca^2^ sensor (D1ER) that had been stably expressed. Data are shown in Figure 10. Ionomycin, thapsigargin and 100 μM amlodipine all evoked prominent ER Ca^2+^ release. 30 μM amlodipine evoked significant Ca^2+^ release and 20 μM amlodipine elicited slow and modest Ca^2+^ release that was nevertheless detectable.

**Figure 10.**
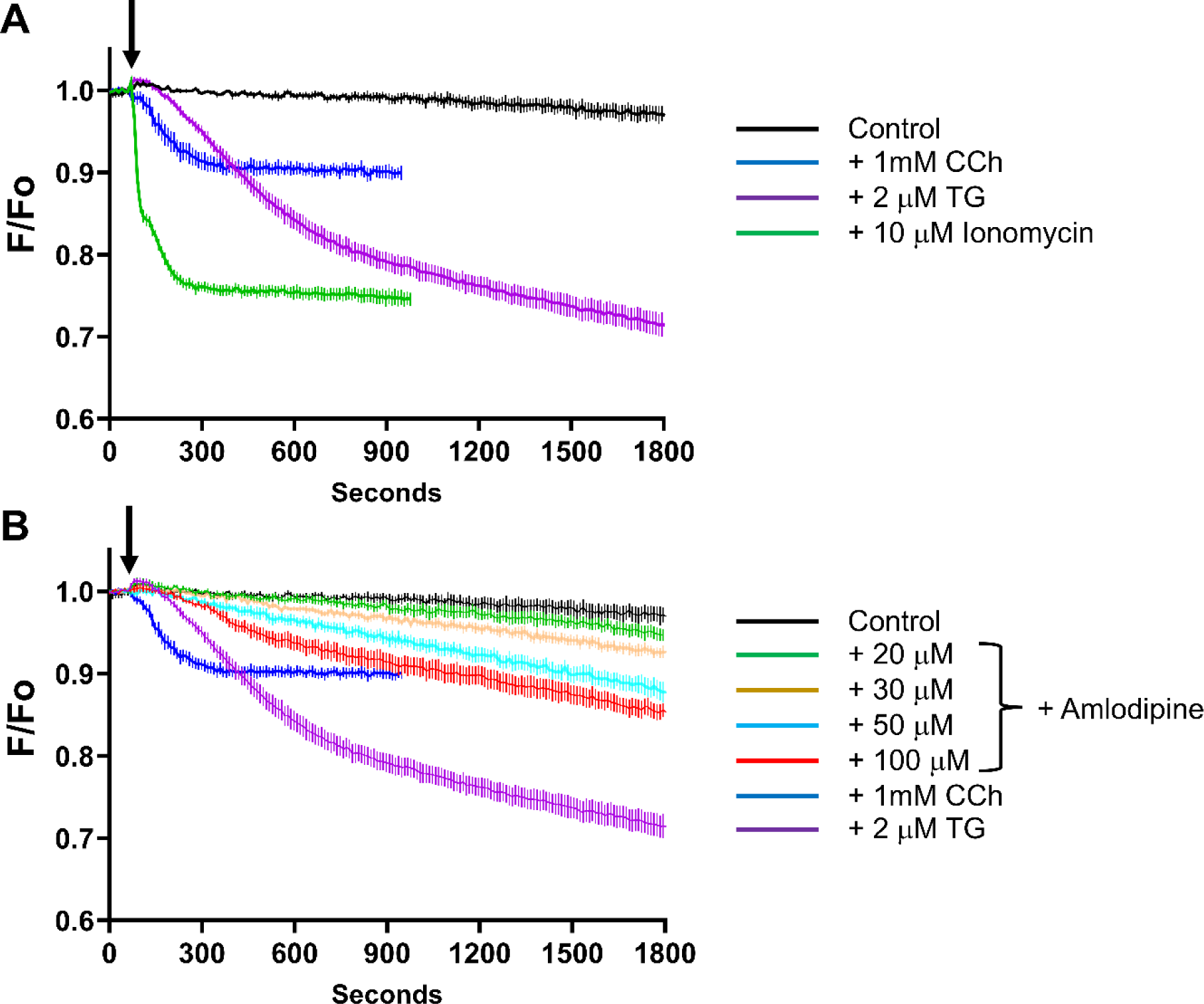
Direct measurement of ER Ca^2+^ (Ca^2+^) using the ER-targeted Ca^2+^-sensitive probe D1ER cameleon. A, Validation of the measurements; ionomycin (green trace), thapsigargin (purple trace) and carbachol (blue trace) all decrease Ca^2+^. B, amlodipine dose-dependently decreases Ca^2+^ at concentrations ≥ 20 μM. All data are mean ± SEM for n=6 experiments.

There are several reasons why our direct ER Ca^2+^ measurements differ from Johnson et al^7^ First, they transiently transfected GCaMP6-150, which would lead to variability between cells and reduce bandwidth of the recordings in cells with relatively low levels of ER-targeted probe. This would be overlooked because the data are presented as F/F0 (Figure 4D of Johnson et al. ^7^). Second, there is enormous variability in their experiments, for each condition^7^. In Figure 4E of Johnson et al. tremendous spread is seen, where a sizeable fraction of cells shows significant store depletion to vehicle and amlodipine, and other cells show significant Ca^2+^ loading of the ER of >30% resting levels under the same conditions.^7^ A supra-maximal concentration of carbachol reduced store content by ∼40%^7^, so many cells show a substantial ER Ca^2+^ decrease both in vehicle and amlodipine (Figure 4E of Johnson et al^7^). Such marked decreases and increases in ER Ca^2+^ above resting levels to the same stimulus (amlodipine) or no stimulus would mask a small decrease in ER Ca^2+^ to 20 μM amlodipine. We also note that the above experiments were conducted in 2mM Ca^2+^-containing external solution^7^, which would allow store refilling following partial ER Ca^2+^ release by amlodipine, especially if a small peripheral and therefore hard to detect sub-compartment of ER is linked to store-operated Ca^2+^ entry, as suggested by Dr Machaca^39^.

## Conclusion

We maintain that the Johnson et al. study has fundamental shortcomings in methodology, lacks several key control experiments that are essential for interpreting the data, contradicts previous recent work from the same group and draws conclusions that are not supported by their own data. Furthermore, the critical analyses of patients’ records, which formed the basis of the claim that LCCBs increase heart failure more than any other intervention, suffers from major mathematical misunderstandings, errors, wrong calculations and lack of consideration of any of a multitude of confounding factors.

The response by Dr Trebak and colleagues has done little to dispel any of our concerns, as the new experiments we have undertaken demonstrate. We therefore stand by the original conclusions of our previous work.

## Acknowledgements

This work was supported by the US National Institutes of Health Intramural Research Program; US National Institute of Environmental Health Sciences (NIEHS) (ZIA ES103353-03 to A.B.P.). We are grateful to Flow Cytometry and Fluorescence Imaging & Microscopy Centres at NIEHS for help with experiments, and the technical assistance of Mr. Scott Gabel and Dr. Eugene DeRose in the NMR Research Core Facility.

